# Genomic signature of shifts in selection in a sub-alpine ant and its physiological adaptations

**DOI:** 10.1101/696948

**Authors:** Francesco Cicconardi, Patrick Krapf, Ilda D’Annessa, Alexander Gamisch, Herbert C Wagner, Andrew D Nguyen, Evan P Economo, Alexander S Mikheyev, Benoit Guénard, Reingard Grabherr, Wolfgang Arthofer, Daniele di Marino, Florian M Steiner, Birgit C Schlick-Steiner

## Abstract

Understanding how organisms adapt to extreme environments is fundamental and can provide insightful case studies for both evolutionary biology and climate-change biology. Here, we take advantage of the vast diversity of lifestyles in ants to identify genomic signatures of adaptation to extreme habitats such as high altitude. We hypothesised two parallel patterns would occur in a genome adapting to an extreme habitat: *i*) strong positive selection on genes related to adaptation and, *ii*) a relaxation of previous purifying selection. We tested this hypothesis by sequencing the high-elevation specialist *Tetramorium alpestre* and four other phylogenetic related species. In support of our hypothesis, we recorded a strong shift of selective forces in *T. alpestre*, in particular a stronger magnitude of diversifying and relaxed selection when compared to all other ants. We further disentangled candidate molecular adaptations in both gene expression and protein-coding sequence that were identified by our genome-wide analyses. In particular, we demonstrate that *T. alpestre* has *i*) a derived level of expression for *stv* and other heat-shock proteins in chill shock tests, and *ii*) enzymatic enhancement of Hex-T1, a rate-limiting regulatory enzyme that controls the entry of glucose into the glycolytic pathway. Together, our analyses highlight the adaptive molecular changes that support colonisation of high-altitude environments.

## Introduction

Adaptation of organisms to climate drives variation in diversification rates and species richness among clades. Their climatic niche affects a species’ distribution in space and time (Soberón 2007), critically affecting both speciation and extinction. Therefore, understanding how organisms manage thermal adaptation is important in terms of both evolutionary biology and climate-change biology, considering the potential world-wide loss of ecological niches (Lamprecht et al. 2018; Rogora et al. 2018). In this context, high elevations, characterized by a short growing season and low annual minimum and mean temperatures, with high daily fluctuation temperatures (Körner et al. 2011), are an important open-air laboratory to study speciation and adaptation to cold habitats; and compared to the great effort in gene-based studies on cold tolerance of model organisms (Clark and Worland 2008), a lesser effort has been put towards understanding truly cold-tolerant animals at the genomic level (Clark and Worland 2008; Parker et al. 2018). This is even more evident when considering that among the approximately 600 sequenced insect genomes available today only four belong to species ecologically restricted to high altitudes or Antarctic habitats (Keeling et al. 2013; Kelley et al. 2014; Macdonald et al. 2016; Cicconardi, Di Marino, et al. 2017).

As yet, potential patterns of genomic signatures in arthropods adapting to more extreme habitats, such as high elevations, have been scarse and, more generally, there is no theory that predicts the rates of genomic change for extreme habitats. Here, we hypothesise two parallel patterns to occur in a genome adapting to an extreme habitat. (1) Strong positive selection on genes related to adaptation, such as genes involved in metabolic pathways; and (2) Relaxation of previous selecting forces, as the conditions of the previous niche are lacking in the new niche. The latter should lead to a reduced number and/or a different set of genes under purifying selection. Specifically, in the case of high-elevation habitats, some heat-shock proteins (HSPs), necessary for coping with extreme heat, should be under relaxation, because heat resistance is not strongly selected for in the average alpine species. A problem with the hypothesis of these two parallel patterns is that there is no direct test available for the functional consequences of the loss or gain of a specific gene. There are often many pleiotropic effects for genes that make interpretations difficult, and HSPs, expressed upon exposure to stress or during development and growth, are likely to fall into this category (King and MacRae 2015). Therefore, we expect strong positive selection in genes involved in energetic metabolism, and an increase in the relaxation rates in other genes, without being able to specify which these might be. Identifying such genes will help to set up hypotheses to test in the future. A comparative approach offers the strongest method for testing our hypothesis, by comparing genomes of species closely related to each other, but divergent in terms of adaptation to different environments. However, this approach can be limited by insufficient niche divergence within a group or the number of genomes sequenced. Ants are emerging as a leading system for comparative genomics due to the constantly increasing number of available genomes. This resource offers the opportunity to improve the accuracy of orthology detection (Nygaard et al. 2016), to scan for specific mutations, candidate genes, and patterns of acceleration and relaxation in the ant genomes associated with adaptation to cold habitats.

Ants are key species in the Earth’s terrestrial ecosystems. They comprise more than 15,442 described species and display an impressive diversity of lifestyles (Hölldobler and Wilson 1990; Bolton 2018; Seifert 2018). Ants are especially notable among insects for their ecological dominance as predators, scavengers, and indirect herbivores. They compose at least one third of the entire insect biomass (Wilson 1990) and colonise all kinds of habitats, including thermobiologically challenging environments. The formicine ant *Melophorus bagoti*, for instance, is active during the hottest periods of the summer day, when air temperatures at ant height exceed 50ºC (Christian and Morton 1992). On the other hand, for example, the myrmicine ant *Tetramorium alpestre* inhabits the montane and subalpine belt of the Central and South European mountain systems, with the Alps as its main distribution area (Steiner et al. 2010; Wagner et al. 2017). This species lives mainly between 1300 and 2300 m above sea level (a.s.l.), forages below the ground, and nests are established in cool grassland under stones, in moss, rootage, and dead wood, especially subalpine and alpine grass mats (Seifert 2018). *Tetramorium alpestre* is thus a system well suited for studying both genomic adaptation to high elevation and socio-behavioural evolution.

Here, we tested our hypothesis of both increased diversifying and reduced purifying (relaxing) selection resulting from adaptation to an extreme niche using *T. alpestre*. We newly sequenced its genome and those of four related *Tetramorium* species with diverging ecological niches: *T. immigrans*, a species of the *T. caespitum* complex to which *T. alpestre* also belongs to, in sympatry with *T. alpestre*, but separated by their altitudinal and ecological habitats; *T. parvispinum*, restricted to mountain forest habitats of the Austral-Asian and Indo-Malayan subregions (Liu et al. 2015); and *T. bicarinatum* and *T. simillimum*, invasive generalist species which occur in warmer habitats (Bertelsmeier et al. 2017; Guénard et al. 2017). Our goals were to define the ecological niche of these ant species based on environmental data, and subsequently to perform gene family expansion/contraction and protein-coding scans for signatures of diversifying and relaxing selection. We also annotated the five HSP subfamilies for 19 ant species to test for possible shifts in selection acting on these gene families, and we performed two experiments in *T. alpestre* and its relative, *T. immigrans*: we assessed (i) chill-shock triggered gene-expression patterns of *starvin* (*stv*), a modulator the activity of Hsp70 chaperone machinery during recovery from cold stress, and certain HSPs involved in recovering from extreme cold, and (ii) the temperature dependence of the enzyme activity of Hexokinase type 1 (Hex-T1), a rate-limiting and regulatory enzyme that controls the entry of glucose into the glycolytic pathway, one of the most conserved and essential hexokinase isoenzymes (Jayakumar et al. 2007). The genomic data presented here contribute to a foundation for studying insect genome evolution, particularly also in the light of climate change.

## Results

### Climatic niches

To define the environment of *T. alpestre* and the other four *Tetramorium* species, a dataset contains 1835 localities representing 17 species was assembled, using georeferenced occurrence data compiled from the Global Ant Biodiversity Informatics database (see method section), with the number of localities per species ranging from 9 to 591 (Table S1). Of the 19 bioclimatic variables, four (bio1: Annual Mean Temperature, bio5: Max Temperature of Warmest Month, bio8: Mean Temperature of Wettest Quarter, bio10: Mean Temperature of Warmest Quarter) differentiated between *T. alpestre* and other ants (adjusted *P*-values < 0.001) (Figure 1c). In detail, *T. immigrans* and *T. alpestre* occupy habitats that are colder in the growing season than those of the remaining species, with *T. alpestre* revealing even colder habitats compared with *T. immigrans*.

**Figure 1.**
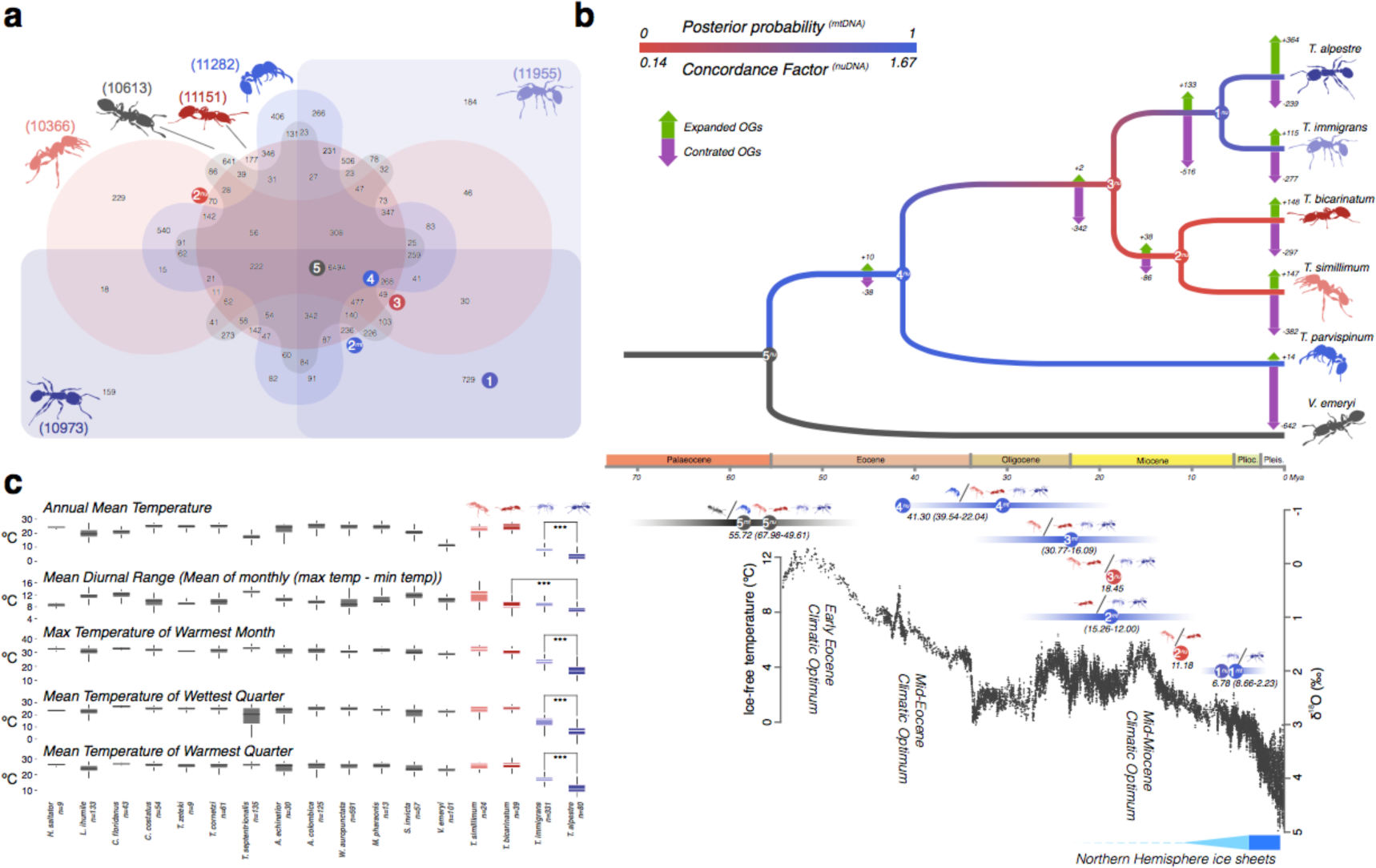
*a*) Venn diagram displaying overlap in orthologous genes in the five *Tetramorium* spp. + *V. emeryi* (Crematogastrini). Numbers close to the species diagrams represent the total number of ortholog groups. Coloured circles with number represent phylogenetic splits in the nuDNA (N^nu^) and mitochondrial phylogenies in *b*; *b*) The dated Crematogastrini nuclear phylogeny (above). Branch colour are based on the concordance factors of the two datasets (nuDNA and mtDNA). On each branch, the numbers of expanded (green arrows) or contracted (purple arrows) genes is shown, inferred from observed OGs sizes at terminal branches. In the bottom, the global δ^18^O (‰) derived from analyses of two common and long-lived benthic taxa are given, *Cibicidoides* and *Nuttallides*, which reflect the global deep-sea oxygen and carbon isotope and thus the temperature (from Zachos et al., 2001). Bars represent the 95% confidence interval for the Bayesian Inference analysis, circles with numbers the phylogenetic splits in the nuDNA (N^nu^) and mtDNA (N^mt^) phylogenies. *c*) Boxplots of the five bioclimatic variables significantly different for *T. alpestre* and the other ant species.

### Sequencing, assembly, and annotation of the *Tetramorium* genomes

The combination of overlapping paired-end libraries with very high coverage, standard short paired-end libraries, and mate pair libraries resulted in an overall complete assembly of the five *de novo Tetramorium* genomes, at the level of both contiguity and scaffolding. In particular, the *T. alpestre* genome contiguity (contig N50) is the 3^rd^ best among all ant genomes available (Table S1). The analysis of the *k*-mer frequency distribution generated was unimodal (Figure S1; Supplementary Material online), and an analysis of the distribution of GC content across the five *de novo Tetramorium* genomes revealed similar distributions with a mean of 38%, that is, slightly less than that found in *V. emeryi* (42%) (Figure S2). No significant bias was recovered considering GC content and coverage (Figure S3). All five *de novo* draft assemblies were ∼241 Mb in length, comparable in size with the average of all ant genomes available (∼278 Mb) (Table S1). The total *T. alpestre* genome size was 245.72 Mb, including 18 Mb of gaps and unknown characters (N/X), slightly smaller than the estimated genome size by flow-cytometry of 291.84 Mb +/-1.76 Mb (n=12). Repetitive elements made up similar proportions of each sequenced genome (*T. alpestre* = 18%, *T. immigrans* = 20%, *T. bicarinatum* = 21%, *T. simillimum* = 20%, *T. parvispinum* = 23%; Table S1), indicating a strong correlation between genome size and total interspersed repeat content (Pearson *ρ* = 0.976, *P*-value < 0.005; Figure S5). For the gene annotations in *T. alpestre*, a combinatorial approach of unsupervised RNA-seq-based, homology-based, *ab initio*, and finally *de novo* methods were used. By doing so, we approached the number of predicted protein-coding genes in the phylogenetically closest annotated species (15,085 genes predicted vs. 14,872 in the *V. emeryi* genome, Table S1). This improved the homology-based annotation of the remaining four *Tetramorium* genome assemblies that did not differ much in terms of estimated gene content, ranging from 16k to 15k annotated loci (Table S1). BUSCO analyses estimated overall good representation, with recovered genes in between the 97.6% and 99.9% (Table S1; Figure S6), and all statistics on mRNA, coding regions (CDS), exon, and intron length gave highly similar and overlapping distributions among all *Tetramorium* species and *D. melanogaster* as a reference (Figure S7). Also, the transcript completeness gave good results with a comparable distribution of percentage of alignment across species (Figure S8).

The orthology search analysis produced a total of 7195 OGs between *D. melanogaster* and Hymenoptera; 3261 of these represented scOGs present in all 22 species analysed. Restricting the orthology analysis to *Tetramorium* spp. and *V. emeryi* resulted in 6494 OGs; among *Tetramorium*, the highest fraction was recovered between *T. alpestre* and *T. immigrans* with 729 OGs, followed by *T. simillimum* and *T. parvispinum* (540 OGs) (Figure 1a). For all *Tetramorium* species., the fraction of orthologous genes with at least a match with one of the three outgroups (*A. mellifera*, *N. vitripennis*, *D. melanogaster*) was on average 7286, with the highest in *T. alpestre* (7879) and the lowest in *T. parvispinum* (6977), in line with the average value found in ants (7831).

### Assembly and annotation of ant mitogenomes and their gene translocations

We assembled complete mtDNA genome sequences from 24 ant species. Because of the differences among SRA datasets, we used different numbers of reads to assemble the complete sequences. The average GC-content for coding genes was 23.9 ± 3.0% and varied slightly among taxa with the lowest in *Formica fusca* (19.0%) and the highest in *Leptomyrmex pallens* (32.8%) (Table S1). While almost all species had the typical gene order and orientation, some rearrangements were observed. All myrmicine ants were characterized by a translocation of *trnV* after *rrnS*; *Solenopsis* spp. had a swapped position of *rrnS* and *trnN*; *Monomorium pharaonis* had at least three rearrangements: the inversion of *nad6*, *cob*, and *trnS2*, the translocation of *trnE* upstream to *trnA*, and the translocation of *trnR* upstream to *trnE*; *Tetramorium* spp. underwent a swap of *trnR* and *trnN* (Figure S9).

### Phylogenetic analyses of *Tetramorium* spp., their diversification dating, and the Miocene-Pliocene origin of the lineage leading to *T. alpestre*

The estimated divergence ages from both datasets (mtDNA, nuDNA) were congruent for most nodes, except for recent nodes within the Attini (Figure 1b; Figure S10-13). In this case, the penalized likelihood approach estimates were always more recent than the estimations made with BI using BEAST. The dating of basal nodes like the MRCAs (e.g., Crematogastrini, Myrmicinae, and Formicinae) were congruent with a previous analysis based on more taxa but fewer genes (Moreau and Bell 2013). The two methods also converged regarding the date of the most recent common ancestor (MRCA) between *T. alpestre* and *T. immigrans*. In that specific case, PL dated that split at 6.8 million years ago (mya) within the confidence interval given by BI (Median: 5.15 mya; 95% HPD: 2.66, 8.23).

The phylogenetic relationships were generally highly congruent and with predominantly strong support (Figure 1b and Figures S11-14). In the ML phylogeny of scOGs (nuDNA), all nodes were supported with a bootstrap value of 1. Within the clade of Crematogastrini (*Tetramorium* spp. + *V. emeryi*), the relation of *T. bicarinatum* and *T. simillimum* was uncertain: nuDNA phylogeny recovered them as sister species with low CU (0.11), while mtDNA phylogeny placed *T. bicarinatum* as sister to the *T. alpestre*-*T. immigrans* cluster, with a bootstrap value of 0.77 and a posterior probability of 0.99.

### Contractions of OGs in the *T. immigrans* and *T. alpestre* lineages and expansion of retrotransposon-related genes in *T. alpestre*

We examined the evolutionary dynamics of OGs looking at more dynamic ranges corresponding with terminal branches. We observed an increased number of OG deaths in the *T. parvispinum* lineage (– 642) and in the lineage leading to *T. alpestre* and *T. immigrans* (– 516). Ortholog groups showed very little expansion in internal branches of the *Tetramorium* radiation. Expanded OGs were common in terminal branches, except for *T. parvispinum* with very few expanded OGs (+ 14), suggesting a possible lack of annotation. *Tetramorium alpestre* showed the highest number of expanded OGs (+ 364) (Figure 1b), with more than 370 of them (∼ 40%) belonging to genes with reverse transcriptase (PFAM: rve, rve_1, rve_2) or transposase and retrotransposon activities (PFAM: Retrotrans_gag, Retrotran_gag_2). Among other PFAM, haemolymph juvenile hormone-binding proteins (PFAM: JHBP; 20 genes), zinc fingers (PFAM: zf-CCHC, zf-H2C2, zf-met, 32 genes), and sugar transporters (PFAM: Sugar_tr, 4 genes) were found.

### Dual signature of evolutionary pressures in single-copy OGs in the *T. alpestre* genome

We investigated the impact of evolutionary pressures on one-to-one orthology in the five *Tetramorium* spp. and 14 other ant species and three outgroups (*Apis mellifera*, *Nasonia vitripennis*, and *Drosophila melanogaster*) by computing both mean *ω* (*d_N_/d_S_*) for terminal branches and relaxing selection (*k*) within the Crematogastrini.

The measure of the mean branch *ω* showed an overall high level of purifying selection along the whole ant phylogeny, including for short branches (Roux et al. 2014; Cicconardi, Marcatili, et al. 2017), with median values of 0.10. However, this distribution was significantly shifted in *T. alpestre*, which showed a median value of 0.38 (Wilcoxon rank-sum test ‘*greater*’, *P*-value < 2.2e^-16^), whilst keeping the proportion of *ω* > 1 in each branch unchanged (Figure 2a, S13). The evaluation of *k* in the five *Tetramorium* spp. and *V. emeryi* had a mean of median values of 0.99, that is, twice the value in *T. alpestre* (median *k* = 0.47; Wilcoxon rank-sum test ‘*less*’, *P*-value < 2.2e^-16^) (Figure 2b). We investigated the nature of the increased *ω* (episodic diversifying selection) and decreased *k* (relaxed selection) in *T. alpestre* by combining the two measures. Two evolutionary trajectories appeared to act simultaneously (two tails of the distribution, Figure 2c): One trajectory seems to have led to episodic diversifying selection, promoting the fixation of nonsynonymous mutations with presumably advantageous fitness effects for genes with increased values of *ω* and *k*, also present in the other species. The other trajectory showed a relaxation of the overall purifying selection in genes with increased values of *ω* and decreased values of *k*, only mildly present in *T. immigrans*.

**Figure 2.**
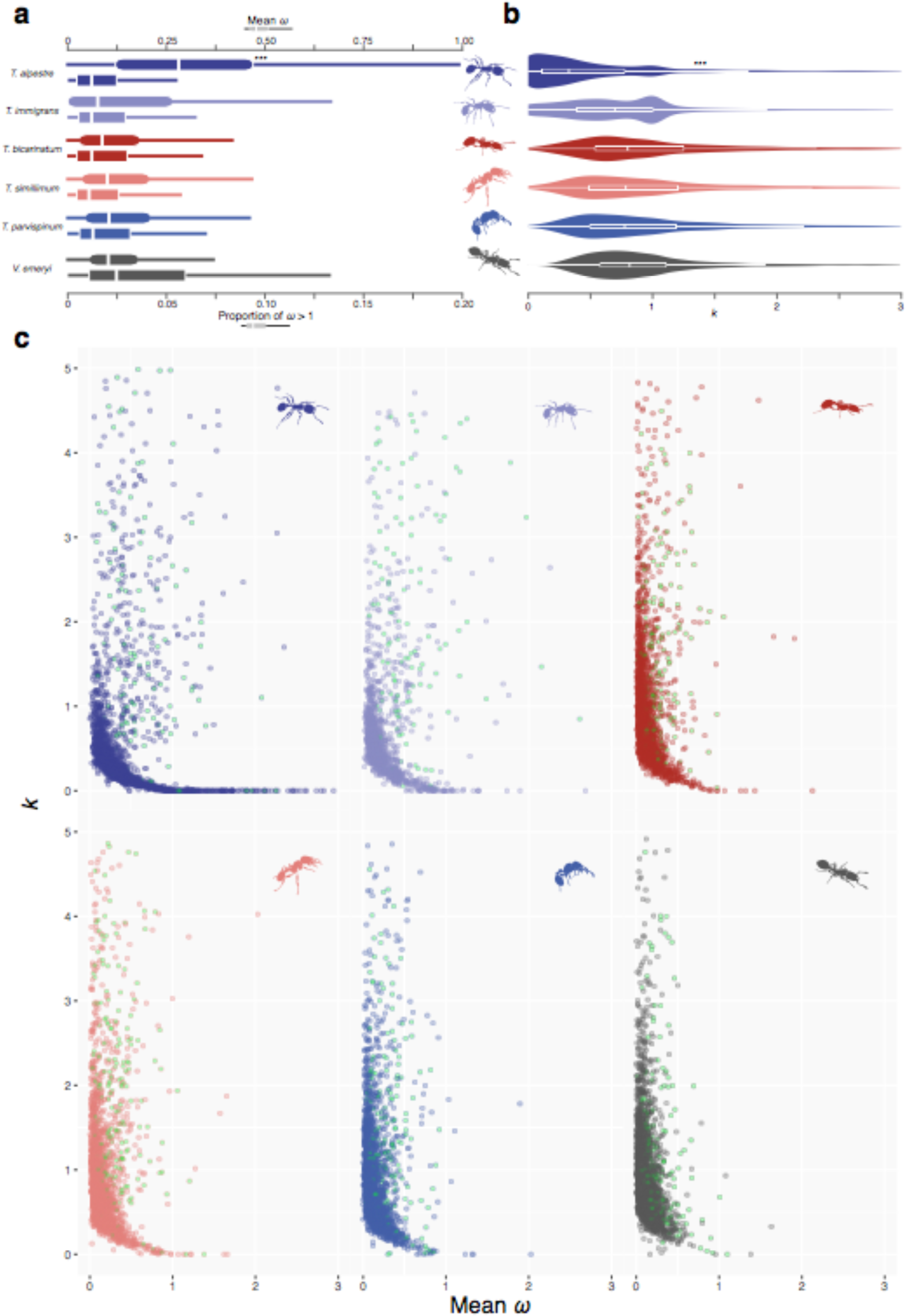
*a)* Boxplots showing the distribution and median (vertical line) of mean *ω* (*d*_N_/*d*_S_) rates (square shapes) in terminal branches of *Tetramorium* spp. and *V. emeryi* (Crematogastrini) in scOGs, and the proportion of genes for which *ω* is higher than one. *b*) Boxplots overlapped by violin plots showing the distribution of *k* in Crematogastrini. The distribution in *T. alpestre* is bimodal as in *T. immigrans*, but with a much more skewed distribution towards zero values. In both sections, asterisks indicate the degree of significance between *T. alpestre* and the other species (Wilcoxon rank-sum tests). *c*) Scatter plots for each Crematogastrini species included, showing for each scOG branch both values of *ω* and *k*. The green dots are scOGs with uncorrected *P*-values associated with diversifying selection less than 0.05. The distribution in *T. alpestre* shows the longest tail corresponding to very low values of *k* and high values of *ω*. It is also notable a much wider and scattered distribution of scOGs with high *k* and *ω* compared with all other species.

After a stringent correction for multiple testing, we found 175 scOGs with a putative signature of diversifying selection (adjusted *P*-values < 0.005; Table S5), enriching 13 biological process terms (*P*-values < 0.005; Figure 3a, Table S6), nine related to cell development (e.g., cell migration involved in gastrulation, photoreceptor cell development, cell fate determination) and organization (e.g., regulation of organelle organization), two related to signalling pathways (enzyme linked receptor protein signalling pathway and torso signalling pathway), one to heart development, and one to organic hydroxy-compound metabolic processes. The most representative genes were *Pten* and *Gbeta13F*, involved in six biological processes, *DCTN1*-*p150* and *dsh*, involved in five of them; all of them involved in the proliferation and division of cells, especially neurological stem cells (*Gbeta13F*), respiratory system development (*Pten* and *dsh*), and the serine/threonine-protein kinase *polo*, present in four enriched Gene Ontology (GO) terms. In the catalytic domain of this protein (255 aa), we found five amino acid substitutions compared with *T. immigrans*, three of them corresponding to active sites (*TalpPolo_G29V_*, *TalpPolo_G30R_*, *TalpPolo_S104R_*). We also tested for possible enrichment in KEGG pathways and found three significant pathways (*P*-values < 0.02; Table S7), all involved in the metabolism of sugars, specifically: glycolysis, pentose phosphate pathway and galactose metabolism. Within these pathways, five enzymes were found under diversifying selection: a glucose-6-phosphate (*CG9008*), the hexokinase type 1 (*Hex-T1*), a phosphofructokinase (*Pfk*), a transketolase (*CG8036*), and a phosphoglucosemutase (*pgm*).

**Figure 3.**
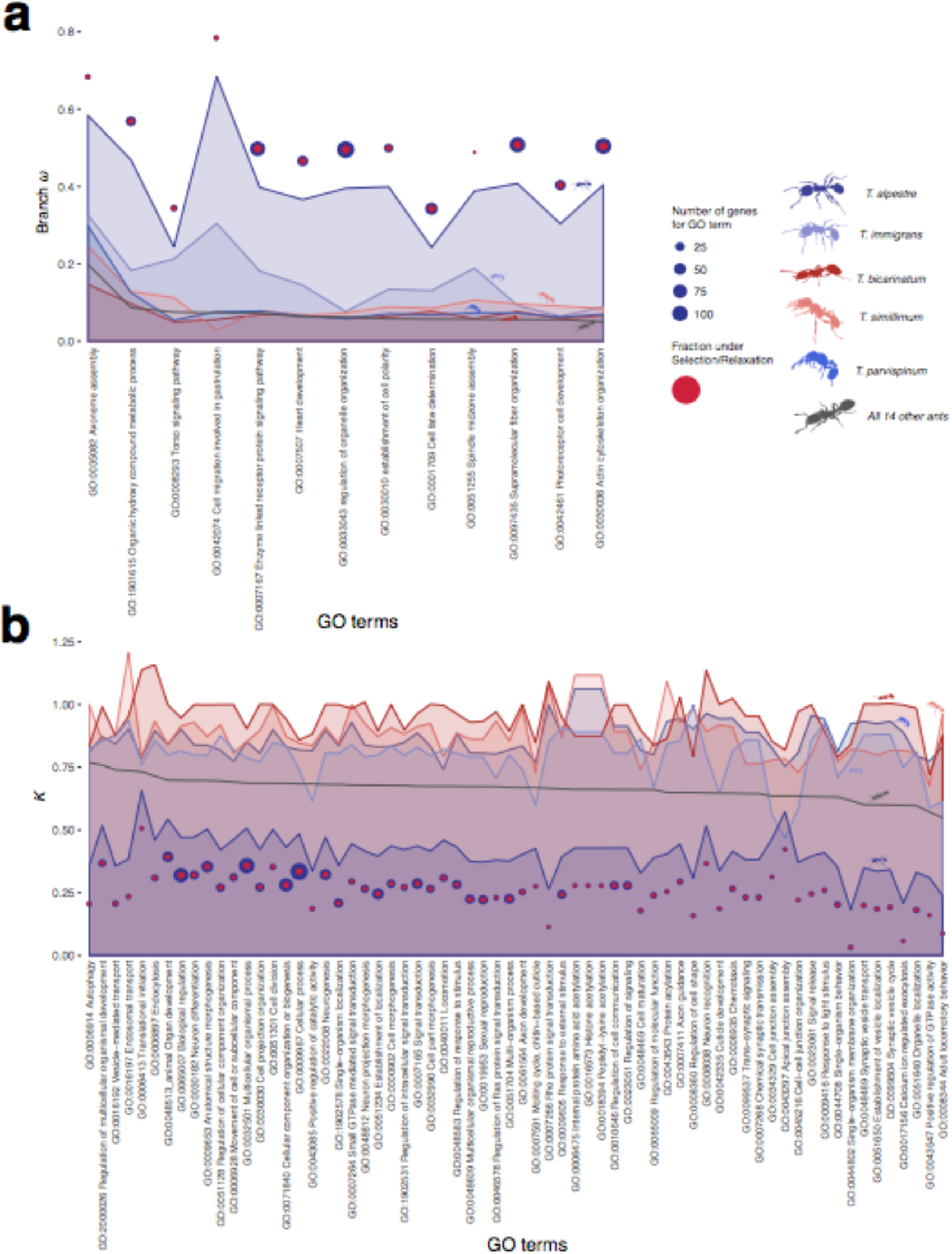
For each enriched Gene Ontology (GO) term category *a*) the median values of *ω*; and *b*) *k* for genes enriching the specific category. *Tetramorium* spp. are colour coded, the other 14 species, merged to obtain a background signal, are in gray. Blue circles are scOGs tested for each GO term, red circles are scOGs returned as significant. Circle sizes are proportional to the number of genes. To be noted are the overall extremely higher values of *ω* and lower values of *k* in *T. alpestre* branches compared to the other species. Seldomly also *T. immigrans* shows few GO terms with high *ω* and low *k* values, in particular the Apical junction assembly (GO:0043297) shows and inverted trend between *T. alpestre* and *T. immigrans*.

132 scOGs had a signature of putative relaxation (*P*-values < 0.005; Table S8). While the absolute number of scOGs under relaxation was slightly lower than the number of genes under diversifying selection, the gene-set enrichment tests revealed many more biological processes significantly enriched: 70 terms (*P*-values < 0.005; Figure 3b, Table S9) enriched by 118 genes. Fifteen terms were related to the regulation of cellular development, signal transduction, and cell communication (e.g., regulation of response to stimuli, regulation of cell communication, regulation of Ras signal transduction), 12 terms related to neurogenesis, axon development, and synaptic transmission (e.g., neuron differentiation, trans-synaptic signalling, axon guidance), seven terms were related to cell morphogenesis and junction organization, and many other related to response to stimuli (e.g., chemotaxis, response to light) or related to adult development and morphogenesis (e.g., anatomical structure morphogenesis, molting cycle, chitin-based cuticle). The enrichment of KEGG pathways revealed three pathways: mRNA surveillance pathway, spliceosome, and protein processing in endoplasmic reticulum (*P*-values < 0.05; Table S10).

### The Hex-T1 activity, its model structure, and molecular dynamics simulations in *T. alpestre* and *T. immigrans*

As we describe in the previous section, the enrichment for KEGG pathways led to the identification of three metabolic pathways, all involved in the metabolism of sugars, with five genes under selection. In order to validate experimentally these *in silico* results, we cloned and expressed the Hex-T1 forms for both *T. alpestre* and *T. immigrans* into *Drosophila melanogaster* cells lines, in order to compare their activities. The purified enzyme of both ants was assayed in at 6, 16, 26, and 36 °C to establish their efficiency under different thermic conditions and identify possible metabolic adaptations to cold-environments. The assays showed that the Km (substrate concentration necessary to reach maximum reaction speed) of Hex-T1 of *T. alpestre* was lower than the enzyme of *T. immigrans* at all temperatures. (Wilcoxon rank-sum test ‘less’, *P*-value < 0.00004, Figure 4a), with values ranging from 0.08 mM to 0.01 mM in *T. alpestre* versus 0.15 mM to 0.05 mM in *T. immigrans*, from the highest to the lowest temperature. These results demonstrate that Hex-T1 of *T. alpestre* was always more efficient than *T. immigrans*, requiring less substrate to reach maximum activity. Although the number of replicates per condition and species was small (n = 3), we performed statistical tests to attempt obtaining insight in the enzymatic activity differences within species at different temperatures and in their variance. Interestingly, it seems that for both species there is an improvement in the Km values shifting towards the lowest temperatures (6 °C *vs* 16 °C; two-way ANOVA followed by a Tukey’s multiple comparison; adjusted *P*-values < 0.05), and a possible deterioration at higher temperatures in *T. alpestre*, from 0.04±0.01 mM (26 °C) to 0.08±0.01 mM (36 °C) (two-way ANOVA followed by a Tukey’s multiple comparison; adjusted *P*-value < 0.0002). An F test (VAR.TEST() implemented in R) to compare the variances of the enzymatic activities (Km values) between the two species – by pooling the replicates at 16 and 26 °C – suggested that there is a possible significant difference in their variance (*P*-value = 0.015), with the confidence interval of Hex-T1 activity in *T. immigrans* twice wider as in *T. alpestre*.

**Figure 4.**
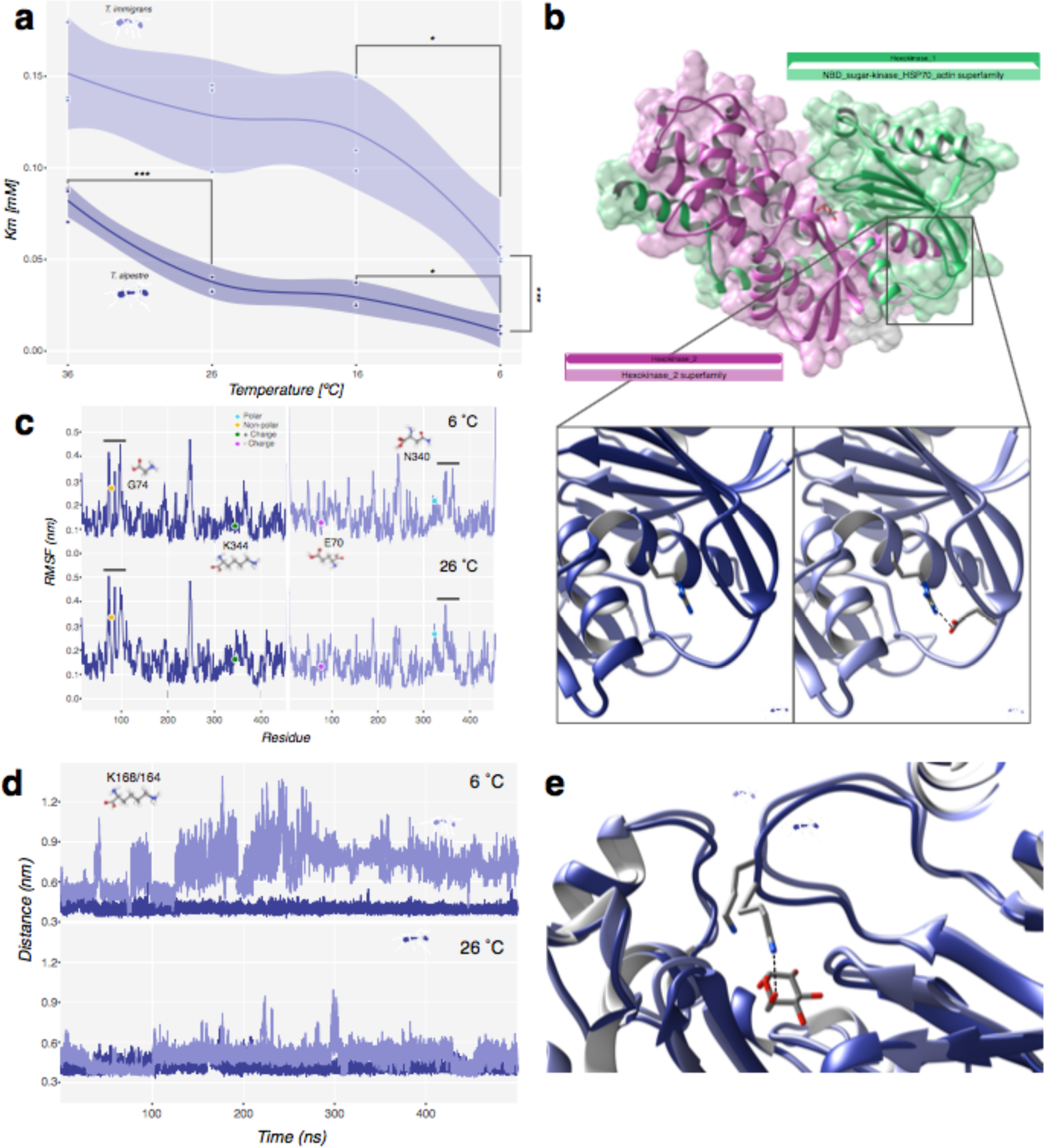
*a*) Scatter plot of Km score across the temperature gradient in the two species, *T. immigrans* on top and *T. alpestre* on the bottom. Lines represent average values with confidence intervals. Significances: *** < 0.001; ** < 0.001; * < 0.05. *b*) Overview of the homology-modeled structure of Hex-T1 in *T. alpestre* showing the two Hexokinase domains, PFAM00349 (green) and PFAM03727 (pink). The lower panels show the location of TalpHex-T1_G74_ (left) and TimmHex-T1_E70_ (right) as well as the arginine residue that forms a salt bridge with E70 (dashed line). *c*) Per-Residue Root Mean Square Fluctuation (RMSF) calculated along 500 ns trajectories carried out at 6 °C (upper panel) and 26 °C (lower panel) for TalpHex-T1 (left panel, blue line) and TimmHex-T1 (right panel, light blue line). The location of the residues found mutated between the two sequences are indicated. *d*) Time evolution of the atomic distance between the centre of mass of glucose and the lateral chain nitrogen atom of TalpHex-T1_K168_ (blue line) and TimmHex-T1_K164_ (light blue line) in the binding pocket calculated along the 500 ns trajectories carried out at 6 °NVTC (upper panel) and 26 °C (lower panel). *e*) Representative configurations extracted from the simulations at 26 °C showing the different orientation of TalpHex-T1_K168_ and TimmHex-T1_K164_, with the first one being the only one able to bind the glucose (dashed line).

The Hex-T1 enzyme is made of two structurally similar domains, Hexokinase 1 and 2 (PFAM00349 and PFAM03727), and an N-terminus that is highly variable both in terms of length and amino acidic composition (Figure 4b). The two domains are relatively well conserved across all ants, other Hymenoptera and *D. melanogaster*. Excluding the hypervariable N-terminus, and considering the two functional domains, the amino acid sequences of *T. alpestre* and *T. immigrans* only differ by two amino-acidic substitutions: a glutamic acid (E) replaced by a glycine (G) in position 74 (TalpHex-T1_E74G_) (Figure 4b inserts) and an asparagine (N) replaced by a lysine (K) in position 344 (TalpHex-T1_N344K_). While the second change probably does not entail a significant functionality effect, the *TalpHex-T1_E74G_* instead may be significant. Because of the lack of the long chain and the negative charge of glutamic acid, this substitution could give the protein a higher flexibility and therefore an overall higher kinetics. We tested this hypothesis by simulating the protein dynamics in the presence of the glucose substrate at different temperatures, i.e. 6 and 26 °C, in the two species, with the intent of mirroring the condition in the in vitro assay. In the simulations we can see a dual effect. For TalpHex-T1_N344K_, located at the C-terminal of a α-helix forming a helix-turn-helix motif (VSETE**K**DPKG), resulted in a lower degree of Root Mean Square Fluctuation (RMSF) revealed a lower degree of fluctuation in TalpHex-T1 (Figure 4c). This is possibly due to the alternation of positive and negative charges creates a salt-bridges network that stabilizes the whole motif (data not shown). In particular TalpHex-T1_K344_ is involved in a stable interaction with TalpHex-T1_E341_ (data not shown), while TimmHex-T1_N340_ is not available for the formation of salt bridges and mainly establishes hydrogen bonds with TimmHex-T1_S336_ or TimmHex-T1_E337_ at 6 or 26 °C, respectively. Although the *T. alpestre* salt bridge brings a higher flexibility of the helix-turn-helix motive, it does not interfere with the catalytic site, given that this motif is far away in the C-terminal region of the protein. In contrast, at the other region (TalpHex-T1_E74G_), the mutation-caused amino acid change allows the loop to explore a broader conformational space equipping it with a higher degree of flexibility already at 6 °C (Figure 4c). Nevertheless, by increasing the temperature, TalpHex-T1 was subjected to an even higher degree of fluctuation, once again confirming the intrinsic flexibility led by the glycine. Notably, the increased flexibility is also transferred to residues relatively far from the mutation, in particular in regions near the active site (168-169), causing a different pattern of interaction with the substrate (Figure 4d, e). Indeed, TalpHex-T1 established a stronger interaction with glucose at both temperatures with respect to TimmHex-T1 (Figure 4e).

### The heat-shock proteins in ants and their evolution in *T. alpestre*

Members of the five subfamilies of the heat shock proteins (HSPs: Hsp90s, Hsp70s, Hsp60s, Hsp40s, and sHsps) were fully recovered and characterized in 19 ant species. Overall, a strong purifying selection was observed within subfamilies (Figure 5a, Table S11). In the Hsp90 gene family (*trap1*, *gp93*, and *Hsp83*) *Hsp83* was found duplicated within the lineage of *W. auropunctata* (inparalog), as well as the *T. parvispinum gp93*; duplication that probably occurred before emergence of the lineage (outparalog) (Figure S14). Members of the Hsp70 family were recovered for all species with 11 ortholog groups, with few duplications in *Hsc70-3* and *Hsc70-5* (two inparalogs and two outparalogs, respectively). We confirmed the lack of Hsp70s in ants (Nguyen et al. 2016) and a multiple duplication of *Hsc70-4* in Hymenoptera. One paralog seemed to be lost in many species while the other two copies were still present in almost all species, with the presence of other outparalogs in various species, mostly in Attini (Figure S15). The chaperonin Cpn60/TCP-1 family (Hsp60) had nine OGs; six inparalogs were found in *CCT7, CCT3* and four outparalogs in *Tcp1a*, *CCT8*. The heat shock protein 40 kDa/Chaperone DnaJ was the largest subfamily, with 37 well-defined OGs. Its turnover of duplication was relatively small, and only six OGs showed some level of duplication. The small HSPs (sHsps) showed the expansion of the lethal (2) essential for life (*l(2)efl*) gene paralog of *D. melanogaster*, with five distinct OGs in all Hymenoptera. *L(2)efl3* seemed to be retained by only in 11 species, and with *l(2)efl4*, they are the two most divergent OGs, with *ω* equal to 0.23 and 0.33, respectively. The less divergent were *Hspb1* and *l(2)efl5*, for which *ω* was 0.11 and 0.07, respectively. Overall, the sHsps showed a significantly higher distribution of *ω* (median = 0.22, *P*-value = 0.013), followed by Hsp40s, which also showed a wide distribution of *ω*. In contrast, Hsp90s, Hsp70s, and Hsp60s seemed to be very stable and under a very high purifying selection.

**Figure 5.**
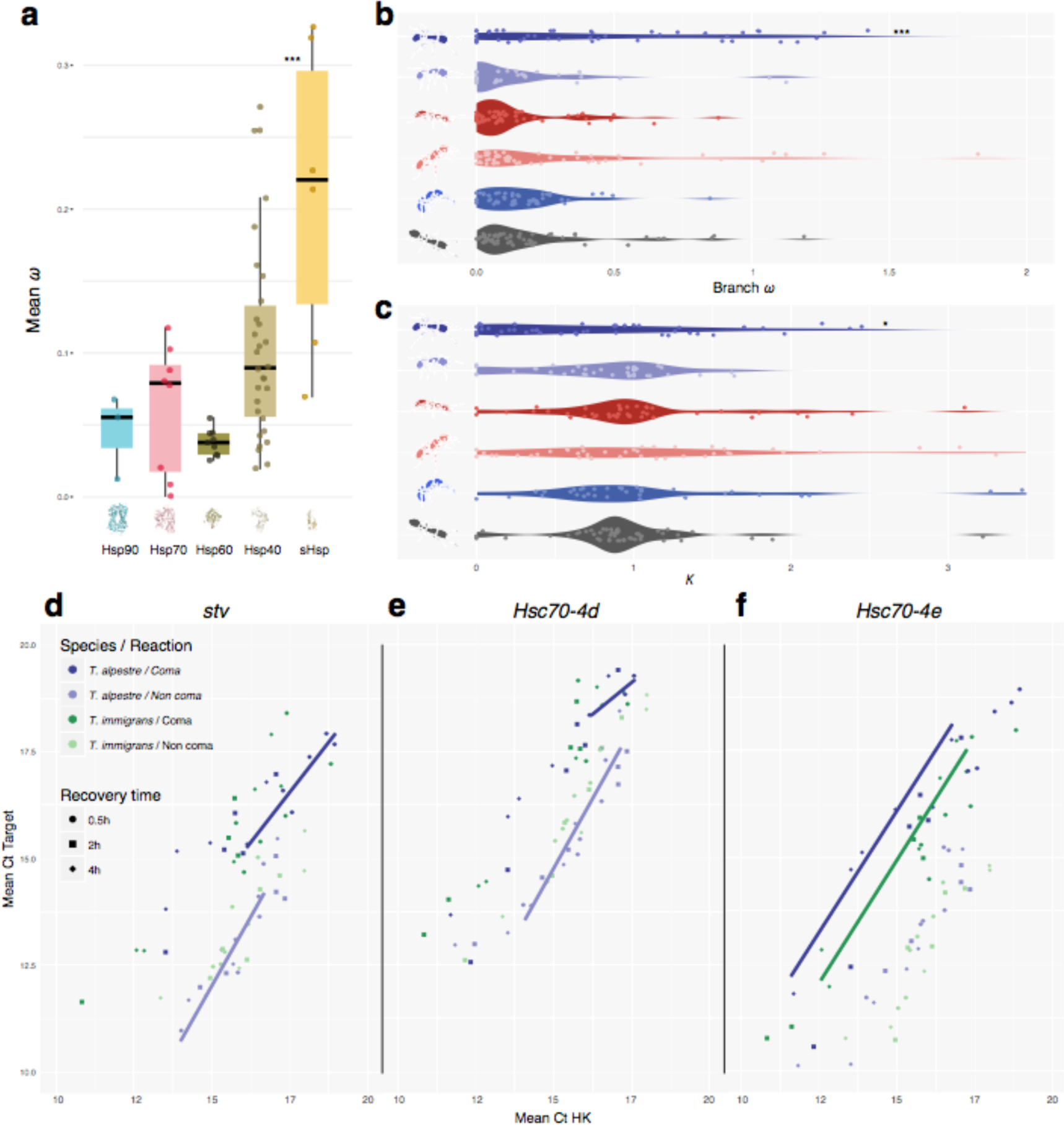
*a*) Boxplots showing the distribution and median (horizontal black line) of mean *ω* (*d*_N_/*d*_S_) rates for each of the HSP subfamily. *b*) Violin plot showing the distribution of the mean *ω* values; and *c*) *k* in the terminal branches of the Crematogastrini (from up to bottom: *T. alpestre*, *T. immigrans*, *T. bicarinatum*, *T. simillimum*, *T. parvispinum*, *V. emeryi*). *d-f*) Scatter plot of target gene (*stv*, *Hsc70-4d*, *Hsc70-4e*) vs. housekeeping gene (HK) concentrations; concentrations given as total number of cycles minus cycle threshold (Ct) values, that is, higher values represent higher concentrations. Straight lines are linear regressions for each target gene against HK using the different recovery times separately; based on one-way analysis of covariance followed by correction for multiple comparison, just lines for those treatment / recovery-time combinations are shown for which significances for just a single species arose, that is, *T. alpestre* coma vs. non-coma after 0.5 h (*d*, *e*; no difference in *T. immigrans*) or for which the two species differed, that is, *T. alpestre* coma vs *T. immigrans* coma after 4.0 h (*f*).

To understand other possible mechanisms of adaptation of *T. alpestre* to the cold, for the HSPs we computed the mean values of *ω* and *k* for all branches leading to *Tetramorium* spp. and *V. emeryi*, integrating 13 ant species, and scanned for diversifying and relaxed selection specifically in *T. alpestre*. The first part of the analysis showed a similar landscape as observed in scOGs. Among the five *Tetramorium* species and *V. emeryi*, five species display a distribution of the mean-branch *ω* ranging between 0.07 (*T. immigrans*) and 0.13 (*T. parvispinum*); *T. alpestre* instead showed significant 2.7-fold higher value of the median mean-branch *ω* (*ω* = 0.36, Wilcoxon rank-sum test ‘*greater*’, adjusted *P*-values < 0.004, Figure 5b). The distribution of *k* among HSPs in the six species showed median values of *k* all close to the expected (*k* = 1), between 0.90 in *V. emeryi* and 1.05 in *T. simillimum*, again with the exception of *T. alpestre*, where the median *k* = 0.63, 1.4-fold lower than the lowest median *k* (Wilcoxon rank-sum test ‘*less*’, adjusted *P*-values < 0.05, Figure 5c). The scan for diversifying positive and relaxed selection in branches of the HSPs of *T. alpestre* showed no gene under putative diversifying positive selection, while six loci were identified to be under putative relaxing selection: one Hsp90 (*Trap1*), two Hsp60 (*CCT4*, *CCT5*), and three Hsp40s (*Sec63*, *DnajC8*, *DnajC11*) (*k* < 0.14, adjusted *P*-values < 0.005). Using these genes in prediction interaction networks, we found that the two Hsp60s interacted with other 21 genes, giving 17 significant enriched functions (FDR < 0.05), mainly related to protein folding and microtubule, spindle organization; the three Hsp40s interacted with 21 genes, giving two enriched functions (FDR < 0.05), related to response to heat and response to temperature stimulus (Figure S19).

### Chill shock effect, recovery times, and expression patterns of *stv* and HSPs in *T. alpestre* and *T. immigrans*

Among all the genes with a putative signature of diversifying selection (see previous sections), we also found *starvin* (*stv*), a BAG-family member which seems to be implicated in Hsp70 ATPase activity during cold recovery in *D. melanogaster* (Colinet and Hoffmann 2010). Therefore, we explored if the modulation of its expression may play a role in how *T. alpestre* differently withstands the cold in comparison to the related species *T. immigrans.* We thus performed a pilot test measuring the different expression patterns of *stv*, two paralogs of *D. melanogaster Hsc70-4* (*Hsc70-4d* and *Hsc70-4e*), a member of the Hsp90 subfamily (*Hsp83*), and sHsp *l(2)efl4* in the two species.

We measured the transcription activity of three recovery times (0.5 h, 2.0 h, and 4.0 h) and compared coma and non-coma individuals, within and between species, and found a significant difference in the expression patterns of *stv*, *Hsc70-4d*, *Hsc70-4e*, and *Hsp83*, but not of *l(2)efl4* (one-way ANCOVA followed by Bonferroni-Holm correction). In particular, looking at the different recovery times, *stv* and *Hsc70-4d* responded quickly in *T. alpestre* but not in *T. immigrans,* being significantly expressed only in workers of *T. alpestre* after 0.5 h recovery, but not in *T. immigrans* (Figure 5d-e). Comparing the expression levels between species, we found higher mRNA levels for *Hsc70-4e* in *T. alpestre* after 4 h recovery (Figure 5f).

## Discussion

*Tetramorium alpestre* represents a critical example of evolutionary adaptation to cold environments such as alpine habitats (Figure 1c). By comparing its genome with other 18 ant species, we shed new light on how *T. aspestre* evolved to colonize sub-alpine habitats and proposed it as a model for studies on arthropods adaptation to cold. We found significant concordance between the nuclear and the mitochondrial dated phylogenies, concerning the dating of the speciation of *T. alpestre*, which seems to have diverged from its closest relative, *T. immigrans*, between 2 and 9 mya. This period overlaps with the beginning of a new climatic zonation of the European continent during the Middle and earliest Late Miocene (Zachos et al. 2001). At the end of this period, a rapid uplift (∼5 mya) fundamentally changed the paleogeographic and topographic setting of central and southern Europe and triggered Alpine glaciation (Kuhlemann 2007). Interestingly, both paleoclimatic and paleogeographic changes co-occurred with the origin of the lineage leading to *T. alpestre* (Figure 1b). In fact, while these glaciations dramatically reshaped global biodiversity patterns, eliminating terrestrial biota from many mid- to high-latitude areas, they may have been the main driver of diversification on highlands by reducing gene flow among populations (Wallis et al. 2016).

The present results support our prediction that two parallel patterns will occur in the genome of species adapting to extreme habitats. The genome of *T. alpestre* appears to be under the influence of two strong evolutionary trajectories: on the one hand, a diversifying selective force, two-fold higher compared with all other ants, and on the other hand, a relaxation of the overall purifying selection present in other ants, identified by the skewed distribution of *k* towards zero. Specifically, the diversifying selective force affects more than a hundred genes in many key biological processes, such as those related to development, cell migration, and gastrulation. Genes under the category of biological processes are also under selection in the genome of the Antarctic midge *Belgica antarctica* (Kelley et al. 2014). In fact, it has been shown in vertebrates that early stages are the most fragile ones, having the highest rate of lethal phenotypes during heat shock and UV irradiation, particularly around gastrulation (Uchida et al. 2018). The same may pertain to genes related to cellular-level organization (e.g., spindle midzone assembly, supramolecular fiber organization, actin cytoskeleton organization), where environmental stress may jeopardise a well working cell division machinery, as SNPs associated to long-term cold-adaptation plasticity were found to be present in genes related to cytoskeletal and membrane structural components in *D. melanogaster* (Gerken et al. 2015). Among 25 genes involved in these biological processes, *polo*, a polo-like kinase which plays a central role as regulator of cell division and is required for several events of mitosis and cytokinesis (Archambault et al. 2015), shows three amino acid substitutions in positions corresponding to active binding sites, and thus could be a good candidate for further studies.

The two evolutionary trajectories were predicted in our hypothesis, and while diversifying selection is somewhat expected in species adapting to a new environmental niche, the magnitude of relaxed selection found is more surprising and never recorded. Although both *ω* and *k* distributions in *T. alpestre* are highly skewed, the absolute number of genes under diversifying and relaxed selection are not markedly greater than other species in similar studies (Harpur et al. 2014; Roux et al. 2014; Cicconardi, Marcatili, et al. 2017). This cannot be directly ascribed to the *type I* error rate because the same rate should be equally randomly present in all the other short or long branches of the phylogeny. Nevertheless, *T. alpestre*, as part of a species complex (Steiner et al. 2010; Wagner et al. 2017), may have more recently diverged from more closely related species not yet adapted to the alpine habitat. Magnitude of this relaxed selection is evidenced by the 70 enriched biological processes, and can be seen as the consequence of a shift and/or decreased magnitude in the purifying selection. The direct implication of this relaxation is not clear, especially its effect on genes and their biological processes, such as the regulation of cellular development, signal transduction, cell communication, neurogenesis, axon development, and many others. If the physiology of cold adaptation utilises different metabolic and developmental strategies to minimise and optimise energy consumption, a new balance needs to be reached. The two patterns here observed (diversifying/relaxing selection) might represent this new shift. Interestingly, *T. immigrans*, which inhabit intermediate environmental conditions (Figure 1c), shows also an intermediate pattern of *k*, with an intriguing bimodal distribution (Figure 2b). Therefore, we can speculate that the highly skewed distributions point to ongoing adaptation to a colder climate, and a relaxation of selective forces present in warmer habitats. To our knowledge, this is the first time that this effect is found in an organism, most likely due to the underrepresentation of genomic studies on alpine and cold adapted species, and, importantly, the lack of application of the RELAXED test in a genome-wise manner.

Examining the effect of those evolutionary trajectories, energetic metabolism seems to be significantly involved in the sub-alpine ant’s adaptation. Several clues are pointing in that direction, such as the more specialized diet underlined by the presence of commensal underground aphids in *T. alpestre* nests but not in those of *T. immigrans* (unpublished data), the expansion of OGs related to sugar transporters, and five enzymes under diversifying selection related to metabolism of sugars. We deeply explored and validated one of these enzymes, Hex-T1, a key regulator and rate-limiting enzyme for energy (sugars) metabolism and reactive oxygen species (ROS) activity in insects (Xian-Wu and Wei-Hua 2016), by contrasting its enzymatic activity between *T. alpestre* and *T. immigrans.* We showed that the mutated form in *T. alpestre* is indeed more efficient, with less substrate needed to reach the highest enzymatic activity, with the protein modeling simulations demonstrating that the single amino acid mutation enhances the catalytic efficiency of the enzyme due to the formation of a more functional network of hydrogen bonds with its substrate. In fact, the path of the hydrogen bonds found in TalpHex-T1 resembled the one depicted by the X-ray solved structure of Hex-T1 from (Mulichak et al. 2002), used as a template to obtain the three-dimensional coordinates of both TimmHex-T1 and TalpHex-T1 (see Methods). Mulichak and colleagues (2000) identified their model residues Lys621 and Asp657 as crucial for the catalytic activity of the protein; the corresponding residues TalpHex-T1_K168_ and TalpHex-T1_D204_ are indeed interacting, together with other residues, with glucose in TalpHex-T1 at both 6 and 26 °C. On the other hand, while the interaction with TalpHex-T1_D204_ is conserved, the one with the lysine is lost in TimmHex-T1 and a looser protein-sugar network is formed, explaining the higher catalytic efficiency of TalpHex-T1 versus that of TimmHex-T1. We can indeed infer that the higher flexibility conferred by TalpHex-T1_G74_ increases the shift of the residue TalpHex-T1_K168_, belonging to the catalytic site (Figure 4d, e), allowing it to move closer and establish a stronger interaction with glucose, possibly contributing to lower the Km of the catalytic reaction. In eco-evolutionary terms all these findings hint at a more intimate relation between *T. alpestre* and the aphids. This relationship between ants and aphids is one of the most studied examples of mutualistic relationships in the animal kingdom. Aphids produce honeydew, and in return ants offer protection from predators, parasitoids, fungal infection and adverse conditions. Honeydew contains monosaccharides (glucose and fructose), disaccharides (maltose, sucrose), and Melezitose, a trisaccharide, that is one of the main sugar synthesized by aphids, and which has been found to attract ants and maintain the ant-aphid mutualism (Fischer and Shingleton 2001). Small amounts of amino-acids, proteins, and lipids are also present (Pringle et al. 2014). Usually, this interaction is above ground level, on plant leaves and branches, but there are cases in which ants are specialized to farm subterranean aphids, just like in *Lasius flavus* (Ivens et al. 2012). *Tetramorium alpestre* may have evolved the same behavior through a coevolution of physiological factors such as palatability, fluid intake rate and digestibility of sugar molecules, improving the ants’ survival by enhancing energy intake, a limiting factor in high-elevation ecosystems.

The complete annotation and characterization of the five HSP subfamilies in 19 ant species show good overall conservation, and strong purifying selection was observed in most subfamilies. The exception are sHsps, which seem to be the most diversified HSP subfamily, with the highest and the widest range of *ω*. It is also the only subfamily that took a different evolutionary path compared with Diptera, having very few orthologs in common with them. Given that sHsps are virtually ubiquitous molecular chaperones that can prevent the irreversible aggregation of denaturing proteins (Haslbeck and Vierling 2015), they may have played a central role in the adaptation of ants to strongly different climates and environments. Although we observe a higher distribution of *ω* in *T. alpestre*, no HSPs show a significant sign of diversifying selection. Rather, six loci show relaxed purifying selection (Hsp90: *Trap1*; Hsp60:*CCT4*, *CCT5*; Hsp40: *Sec63*, *DnajC8*, *DnajC11*). The impact of the TNF receptor-associated protein (Trap1) on cellular bioenergetics of *T. alpestre* may contribute to its adaptation to cold environments by differently modulating cellular metabolism. Given that it functions as a negative regulator of mitochondrial respiration, able to modulate the balance between oxidative phosphorylation and aerobic glycolysis, possibly another form of metabolic adaptation. It has been shown that reduced or absent *Trap1* expression deregulated mitochondrial respiration leading to a high energy state. This state is characterized by accumulation of ATP, tricarboxylic acid cycle intermediates and reactive oxygen species, as well as an increase in fatty acid oxidation (Yoshida et al. 2013). *Trap1* overexpression is associated with increased expression of genes associated with cell proliferation (D. Liu et al. 2010). Since the two Hsp60s are associated to protein folding and cytoplasmic microtubule organization, biological processes also enriched by genes under diversifying selection, this suggests profound changes in the cellular physiology of *T. alpestre*. It is also plausible that adapting to a cold environment will depress selection on genes directly related to severe heat shock. As we have shown, *T. alpestre*’s habitat is characterised by temperatures that rarely exceed 20 °C (median = 16.5 °C; 99% confidence interval = 17.8 °C), although rocky habitats may produce warmer microhabitats. This could mean that this species experienced relaxed selection for HSPs associated with the denaturation to protein folding by heat shock, as suggested by the relaxation of purifying selection on *Cct5* and the three Hsp40s associated with responses to heat and temperature stimulus.

Although we provide a range of evidence that adaptation to cold environments can be related to changes in protein-coding sequences, it has long been postulated that phenotypic divergence between closely related species, such as *T. alpestre* and *T. immigrans*, is primarily driven by quantitative and spatiotemporal changes in gene expression, mediated by alterations in regulatory elements (e.g.: Danko et al., 2018; Prescott et al., 2015). We tested this hypothesis by looking at the expression patterns of *stv* and four other HSPs (*Hsc70-4d*, *Hsc70-4e*, *Hsp83*, *l(2)efl4*), between *T. alpestre* and *T. immigrans*. *Stv* is a member of Bcl-2-associated athanogene (BAG)-family proteins, which interact with Hsc70s and Hsp70s, and can modulate, either positively or negatively, the functions of these chaperones (Doong et al. 2002). It is implied in the recovery from chill shock in *D. melanogaster* during cold recovery (Colinet and Hoffmann 2010; Colinet and Hoffmann 2012). The concerted synthesis of *D. melanogaster stv* and Hsp70s suggests cooperation to offset cold injury, possibly by preserving the folding/degradation and by regulating apoptosis (Colinet and Hoffmann 2010). In our experiment, an acute cold stress (−6°C) was used to knock down individuals. By looking at the differential expression of these genes in coma and non-coma individuals within and between species, we were not only able to show that *stv* may interact with Hsc70s, but also that *T. alpestre* has a quicker response in these genes compared with, *T. immigrans,* the related species not cold adapted. Given these results, which are in line with the selection signature we identified, it is probable that *stv* may play a similar role in the physiology of ants. As we have shown in this study, ants completely lack Hsp70s (Figure S14) and retained only the heat shock cognates (Hsc70s) with mainly two conserved paralogs (*Hsc70-4d* and *Hsc70-4e*), which may have evolved to function as the Dipteran Hsp70s. Although we found an interesting and promising correlation between chill shock, *stv*, and Hsc70s, it will be important to continue exploring other expression patterns in ant physiology to assess whether *stv* has tissue-specific expression patterns, as was found in *D. melanogaster* (Coulson et al. 2005), and to study its expression across different species with different ecological adaptations.

One of the most enduring problems of evolutionary biology is explaining how complex adaptive traits originate (Wagner and Lynch 2010). Although it is widely assumed that new traits arise solely from selection on genetic variation, many researchers ask whether phenotypic plasticity precedes and facilitates adaptation (Levis and Pfennig 2016). This alternative scenario, the plasticity-first hypothesis, states that under novel conditions, environmentally induced variation can be refined by selection and, depending on whether plasticity is favoured, become developmentally canalized through genetic assimilation. Under this scenario, environmentally induced phenotypic change can precede and promote the evolutionary origins of a complex adaptive trait.

Four criteria have been proposed for verifying this hypothesis (Levis and Pfennig 2016): 1) the focal trait needs to be environmentally induced in the ancestral-proxy lineages; 2) cryptic genetic variation will be uncovered when ancestral-proxy lineages experience the derived environment; 3) the focal trait will exhibit evidence of having undergone an evolutionary change in its regulation, in its form, or in both in the derived lineages; and 4) the focal trait will exhibit evidence of having undergone adaptive refinement in the derived lineages. Though it is methodically difficult to prove each point, our findings on the evolution and adaptation of *T. alpestre* to cold appear consistent with plasticity-first evolution. In more detail, *T. immigrans*, the ancestral-proxy lineage, shows considerable plasticity under variable temperature conditions compared to other ant species (Criterion 1, Figure 1c), showing an ability to recover from severe chill-shocks almost as good as the derived lineage (*T. alpestre*), and, unexpectedly, better efficiency of Hex-t1 at lower rather than higher temperatures (Figure 4a). We suggest that the derived lineage, *T. alpestre*, underwent genetic assimilation of the cold response (Criterion 3), followed by evolutionary refinement via changes in regulation, resulting, for example, in *stv* differential expression (Criterion 4). Consistent with evolutionary refinement, the derived lineage has evolved more extreme traits such as an overall improved activity of Hex-T1 with reduced variability.

Our comparative genomic analysis shows how natural selection could trigger complicated patterns of change in genomes, especially protein-coding sequences, both in terms of diversifying selection and relaxation of purifying selection. Many of these changes are in genes that may be associated with aspects of development, either directly or through the associated complex changes in ecology and natural history. Furthermore, this work characterized some candidate genes activities, both *in vitro* and *in silico*, confirming our hypothesis and opening for the identification of other genes possibly playing important role in adaptation to new ecological niches. Overall, this study represents the first systematic attempt to provide a framework for the genomic analysis of adaptation to extreme environments and underlines the importance of studying organisms with different ecological niches in understanding the genetic basis of ecological adaptation.

## Methods

### Species-locality and climatic data

To define the environment of *T. alpestre* and the other four *Tetramorium* species, georeferenced occurrence data were compiled and complemented with those for the native ranges of other ant species with available genomic data. Occurrence records for 17 species (Table S1) were compiled from the Global Ant Biodiversity Informatics database (GABI, Guénard et al., 2017, accessible via www.antmaps.org, Janicki et al., 2016). The species determinations in the records were evaluated by specialists in ant taxonomy, and erroneous data were excluded. Slightly incorrect coordinates (i.e. in the ocean within 8.5 km of land) were assigned to the nearest land point. Additionally, georeferenced occurrence data of two species (*T. alpestre, T. immigrans*) were compiled from the latest taxonomic revision (Wagner et al. 2017). The occurrence data of each species were spatially rarefied at a 5 km level using the R package SPTHIN v. 0.1.0 (Aiello-Lammens et al. 2015), to counter spatial autocorrelation and to provide some protection against sampling bias issues (Merow et al. 2013). The Bioclim variables bio1-bio19 from the WORLDCLIM database v. 1.4 (Hijmans et al. 2005) were extracted for each unique species locality at a 2.5 arc min resolution (4.65 x 4.65 = 21.6225 km^2^ at the equator) using ARCGIS v. 10.4 (ESRI, Redland, CA). A Wilcoxon-Mann-Whitney rank sum test as implemented in the R function WILCOX.TEST (http://www.r-project.org) v. 3 (R Development Core Team 2017) was adopted to test for differences between *T. alpestre* and the other species individually for each of the Bioclim variables. The Bonferroni-Holm sequential rejection procedure (Abdi 2010) was used to control for multiple testing.

### Biological sample processing, genome size estimation, and genome sequencing

Six colonies of *T. alpestre* ants were sampled, two for flow cytometry (nest 18811: Penser Joch, Italy, 46.83379° N, 11.44652° E, 21 June 2016; nest 18813: Kühtai, Austria, 47.27923, 11.07590° E, 21 June 2016), three for RNA extraction (nest 18586: Zirmbachalm, Austria, 47.21920° N, 11.07530° E, 30 July 2015; nest 18590: Zirmbachalm, 47.21910° N, 11.07620° E, 30 July 2015; nest 18594: Penser Joch, 46.82940°N, 11.43830° E, 2 August 2015), and one for DNA extraction (nest 18592: Penser Joch, 46.83380° N, 11.44610° E, 2 August 2015). The ants were brought alive to the laboratory (for flow cytometry) or immediately submerged in RNAlater (for RNA extraction) or 96% ethanol (for DNA extraction) and stored at −70 °C (RNA) or −20 °C (DNA).

Flow cytometry was used to estimate relative genome size from single *T. alpestre* worker heads, using strain ISO-1 of *Drosophila melanogaster* as internal standard (obtained from Bloomington Drosophila Stock Center, Indiana University, Bloomington, USA; nuclear DNA content: 0.35 pg, Gregory 2018). Six individuals each were measured from *T. alpestre* nests 18811 and 18813. In more detail, single *T. alpestre* heads and, separately, heads of 10 ISO-1 females were chopped in 500 µL of ice-cold Otto 1 Buffer (0.1 mol/L citric acid, 0.5% Tween20 (Merck KGaA, Darmstadt, Germany)). The suspensions were filtered using 42-µm nylon meshes and incubated for 5 min. Then, 1 mL of staining solution (DAPI (4 µL/mL) and 2-mercaptoethanol (2 µL/mL) in Otto 2 buffer (0.4 mol/L Na_2_HPO_4_·12 H2O)) was added to each suspension. For each measurement, 200 µL suspension of a single *T. alpestre* worker and 10 µL of the ISO-1 standard were used. Fluorescence intensity of 3000 cell nuclei per sample was measured with a Partec CyFlow space flow cytometer (Partec GmbH, Münster, Germany). Gating and peak analysis were done automatically using the software Partec Flo Max.

For the DNA and RNA extractions, the individuals were thawed on ice and the gasters removed. DNA was extracted using the QIAamp DNA Micro Kit (Qiagen). The extraction followed the protocol of the manufacturer, except that 20 µl RNase A solution (Sigma-Aldrich) was added to the vials and incubated for 5 minutes at room temperature. DNA concentration and quality were checked via Nanodrop (Thermo Scientific) and gel electrophoresis. Two adult males were used to generate the overlapping-read and the paired-end (PE) libraries, while for the mate-pair (MP) libraries, four adult females were extracted of nest 18592. RNA was extracted using the RNeasy Mini Kit (Qiagen) following the instructions of the manufacturer. Six developmental stages were extracted as pools of individuals (nest 18586: worker larvae – 30 individuals, worker pupae – 40, worker adult – 50; nest 18594: gyne pupae – 2; nest 18590: gyne adult – 7, male adult – 7). RNA concentration and quality were checked via Nanodrop (Thermo Scientific) and gel electrophoresis.

All *T. alpestre* DNA-seq and RNA-seq Illumina libraries were prepared by IGATech (http://igatechnology.com/). Specifically, four libraries were used for genome sequencing of *T. alpestre*: one overlapping-reads library (insert size ∼400 bp; paired-end (PE) sequencing with 2×250 bp read length), one regular PE library (insert size ∼400 bp, 2×100 bp read length), and two mate pair libraries (insert size ∼4 kb, 2×100 bp read length) (see Table S2). For RNA-seq, six strand-specific libraries (insert size ∼500 bp, 2×200 bp read length) were constructed from three stages of workers (larvae, pupae, adult), two stages of gynes (larvae and adult), and adult males. All libraries were sequenced on an Illumina HiSeq 2500 platform in standard (DNA) and RAPID (RNA) mode. Each of the other *Tetramorium* species were collected kept in EtOH (100%) and identified by Dr. Franscisco Hita Garcia (*T. immigrans*: San Francisco, Golden Gate Park, 37.77035 N°, 122.4688417 W°, 15 September 2014; *T. bicarinatum*: Onna Okinawa Institute of Science and Technology, 26.4622 N°, 127.83056 E°, 14 July 2013; *T. simillimum*: Kibale National Park, Kanyawara Biological Station, 0.56437 N°, 30.36059 E°, 10 August 2010; and *T. parvispinum*: Yunnan Province, Xishuangbanna, XTBG, 21.91625 N°, 101.2739167 E°, 8 June 2013). DNA extraction was performed using the entire body of several workers using DNeasy Blood and Tissue kits (Qiagen), according to the manufacturer’s instructions, and TruSeq libraries (insert size ∼500 bp, 2×100 bp read length) were prepared and sequenced on an Illumina HiSeq 4000 machine at OIST sequencing center.

### Genomic and mitogenomic assembly and gene prediction

Genomic sequencing reads of *T. alpestre* were filtered to remove low-quality reads using TRIMMOMATIC v.0.6 (ILLUMINACLIP:2:30:10 SLIDINGWINDOW:5:15 MINLEN:200) (Bolger et al. 2014), and corrected, and assembled using ALLPATHS-LG v. 52488 (Gnerre et al. 2011). Contigs were first constructed based on overlapping reads and then scaffolded using paired-end and mate-pair information from all DNA libraries. Given the high coverage of the overlapping reads, based on the estimated genome size, multiple trials were performed using different coverages to establish the best combination of read coverage. For the other species TRIMMOMATIC was also used to filter out reads and SPADES v. 3.10.0 (Bankevich et al. 2012) was used to run error correction and the genome assemblies using a *k*mer range of 21, 33, 55, 77, 99, and mismatch correction. For all assemblies, scaffolds belonging to contaminants were identified and removed using BLOBOLOGY v. 2013-09-12 (Kumar et al. 2013).

A bioinformatic pipeline that included combinatorial approaches: homology-based, *ab initio*, and *de novo* methods were implemented to predict gene models for *T. alpestre*. REPEATMASKER v. 4.0.6 (Smit et al. 2013) was used to generate repeat hints. For the homology-based exon hints, protein data sets of *D. melanogaster*, *Apis mellifera*, *Nasonia vitripennis*, and 12 ant species (see Table S1) were aligned to the *T. alpestre* genome using TBLASTN (*e*-value cut-off: 1e^−5^) and EXONERATE v. 2.2.0 (Slater and Birney 2005). As an *ab initio* procedure, the merged RNA-seq libraries were mapped using STAR v. 020201 (Dobin et al. 2013) (settings: ALIGNENDSTYPE Local; ALIGNINTRONMIN 30; ALIGNINTRONMAX 450,000; OUTSJFILTERINTRONMAXVSREADN 80 100 500 1000 2000 5000 20,000; ALIGNSJOVERHANGMIN 10; ALIGNTRANSCRIPTSPERREADNMAX 100,000) to generate the splice-site position-specific scoring matrix (PSSM) and CUFFLINKS v. v2.2.1 (Trapnell et al. 2012) to generate UTR and intron hints. This information was then integrated with GENEMARK-ES v. 4 (Hoff et al. 2015) to train the analysis and use the outcome under the BRAKER1 procedure (Hoff et al. 2015) and AUGUSTUS v. 3.2.1. (Stanke et al. 2006). To avoid missing genes due to unassembled regions of the genome, previously unmapped reads were *de novo* assembled using TRINITY v. 2.2.0 (Grabherr et al. 2011) and annotated with FRAMA (Bens et al. 2016). The three results where then merged, and Benchmarking Universal Single Copy Orthologs (BUSCO v. 1.2) (Simão et al. 2015) was adopted to evaluate the annotation completeness together with integrity and completeness evaluation using DELTABLAST (Boratyn et al. 2012). For *T. alpestre* a visual inspection of the annotation and gene integrity was also performed on more than 250 loci. The generate transcriptome was then used to annotate the other *Tetramorium* species with the other ant proteomes in an homology-based annotation using AUGUSTUS.

All available complete ant mitochondrial genomes (mtDNA) were downloaded from GENBANK (Benson et al. 2014) (Table S1). Illumina raw SRA data of other ant species were downloaded and processed following Cicconardi et al., (2017b). In brief, reads were quality filtered and assembled using IDBA-UD v. 1.1.1 (Peng et al. 2012) and SPADES v. 3.10.0 (Bankevich et al. 2012), and annotated using MITOS v. 2 web server (Bernt et al. 2013).

### Phylogenetic analysis

Nuclear single-copy ortholog genes (scOGs) and complete mitochondrial genomes were used to compute the ant species tree. Once scOGs were determined (see below), single-locus trees and a species tree of concatenated scOGs were estimated using Maximum Likelihood (ML) search as implemented in FASTTREE v. 2.1.8 SSE3 (Price et al. 2010). Gene trees were summarized with MP-EST (L. Liu et al. 2010) using the STRAW web-server (Shaw et al. 2013).

For the mtDNA phylogeny, only protein-coding genes were used. Each gene was aligned using MACSE v. 1.01b (Ranwez et al. 2011), and all alignments were concatenated. The best partitions and models of evolution were identified with PARTITIONFINDER v. 1.1.1_Mac (Lanfear et al. 2014). MtDNA trees were searched with ML and Bayesian Inference (BI) algorithms. For ML, GARLI v. 2.01.1067 (http://code.google.com/p/garli) was used performing 20 + 5 runs from random starting trees. The runs were continued until no further improvement in log-likelihood was found, followed by 1000 bootstrap pseudoreplicates. The results were summarized using SUMTREES v. 3.3.1 (http://bit.ly/DendroPy), using *A. mellifera*, *N. vitripennis*, and *D. melanogaster* as outgroups. For BI, BEAST v. 2 (Bouckaert et al. 2014) was adopted to obtain the dated mtDNA tree and to estimate the divergence of the most recent common ancestors (MRCAs) (template: BEAST (Heled and Drummond 2010)). Fossil data were taken mainly from Moreau & Bell (2013) and PALEOBIO DB (paleobiodb.org) (see Table S4 for a detailed list of references) and used as minimum calibration points constraining nodes in the topology as listed in Table S4. Lognormal prior distribution was implemented with an offset corresponding to the minimum fossil age, log(mean) of 1.0, and log(SD) of 1.0. Ten independent runs of 8 x 10^7^ generations were sampled every 100,000^th^ generation. Each partition was modelled with an uncorrelated relaxed clock with a Yule process, species tree prior, and its best substitution model. Evolutionary models not implemented in BEAUTI v2 (Bouckaert et al. 2014) were manually edited in the xml file. TRACER v. 1.6 (http://beast.bio.ed.ac.uk/Tracer) was used to evaluate convergence and the parameters’ effective sampling size (ESS) and a burn-in was manually implemented and roughly 10% from each run were removed. LOGCOMBINER (Bouckaert et al. 2014) and TREEANNOTATOR (Bouckaert et al. 2014) were used to summarize the results in a single consensus tree. The non-ant species were excluded from the analysis, and no outgroup was adopted. Divergence dates for the maximum-likelihood nuclear DNA (nuDNA) tree (only ants) were also inferred using the penalized likelihood approach (Sanderson 2002) implemented in the R package APE v. 5.1 (Paradis et al. 2004) by calibrating four nodes: the root, the *Tetramorium*, the *Solenopsis*+*Monomorium*, and the Attini branches (Table S4).

### Functional annotation and orthologous-group dynamic evolution

Genes from 19 ant and three outgroup species (Table S3) were clustered into ortholog groups (OGs) with HIERANOID v. 2 (Kaduk and Sonnhammer 2017), which implements INPARANOID v. 8 (Sonnhammer and Östlund 2015) using a guide tree topology based on Moreau & Bell (2013) and BLASTP (*e*-value cutoff of 10^-5^) to improve the reciprocal hit accuracy (Edgar 2010), while homology relationships were searched with DELTABLAST (Boratyn et al. 2012). The functional annotation of each putative protein coding sequence was performed by identifying both the protein domain architecture and Gene Ontology (GO) terms. For each sequence, first HMMER v. 3.1b2 (HMMSCAN) (Eddy 2011) was used to predict PFAM-A v. 31.0 domains, then Domain Annotation by a Multi-objective Approach (DAMA) v. 2 (Bernardes et al. 2015) was applied to identify architectures combining scores of domain matches, previously observed multi-domain co-occurrence, and domain overlapping. Annotation of GO terms was performed as implemented in the CATH assignments for large sequence datasets (Das et al. 2015). Briefly, each input sequence was scanned against the library of CATH functional families (FUNFAMS) HMMs (Sillitoe et al. 2015) using HMMER3 (Eddy 2011) to assign FUNFAMS to regions on the query sequence (with conditional *E*-value < 0.005). Then the GO annotations for a matching FUNFAM were transferred to the query sequence with its confidence scores, calculated by considering the GO term frequency among the annotated sequences. Finally, a non-redundant set of GO annotations was retained, making up the GO annotations for the query protein sequence (Sillitoe et al. 2015).

To identify orthologous groups specifically expanded in the *T. alpestre* genome, BADIRATE v. 1.35 (Librado et al. 2012) was performed twice, once for the nuDNA and one for the mtDNA ultrametric tree topologies, reconstructing ancestral family sizes, gain, death, and innovation (GDI), applying stochastic models and allowing estimation of the family turnover rate (λ) by ML. Using this approach, we observed a correlation between the turnover rate (λ) and branch lengths (Spearman correlation: *ρ* = −0.52, *P*-value = 0), and a bimodal distribution of λ. We interpreted the distribution closer to 0 as background noise and therefore considered only expanded and contracted OGs with λ within the second distribution. The boundary between the two distributions was defined by the valley values between the two, which were computed using OPTIMIZE and APPROXFUN functions implemented in the STATS v. 3.4.0 package in R.

### Heat-shock protein family analysis

From the functional annotation analyses of all Hymenoptera, all sequences bearing a valid protein domain matching HSPs were extracted and aligned using the CLUSTALW v.1.2.1 EBI web server (Larkin et al. 2007) (settings: MBED true; MBEDITERATION true; ITERATIONS 5; GTITERATIONS 5; HMMITERATIONS 5). A phylogenetic tree was computed using ML as implemented in FASTTREE v. 2.1.8 SSE3 (Price et al. 2010) and manually checked to identify and provisionally annotate the five main subfamily proteins (Hsp90, Hsp70, Hsp60, Hsp40, and sHsp) and their OG. Sequences falling outside clusters and with low bootstrap values were manually checked with the DELTABLAST (Boratyn et al. 2012) web-server and with the CONSERVED DOMAIN DATABASE (CDD) web-server (Marchler-Bauer et al. 2017) to identify and remove bacterial contaminants and spurious/incomplete sequences. Then, nucleotide sequences belonging to each subfamily were revers translated into amino acids, aligned using CLUSTALW v.1.2.1, and the obtained protein alignment was used to derive the nucleotide one. FASTTREE was used again to compute phylogenetic trees for the five HSP subfamilies.

### Selection on single-copy orthologous groups and gene families

All selected scOGs and HSP-OGs were scanned to evaluate selection signature on coding regions of *T. alpestre*. This was done by computing the mean *ω* (the ratio of nonsynonymous to synonymous substitution rates; *d*_N_/*d*_S_) and the relaxation of each branch of the phylogeny of *Tetramorium* spp. + *Vollenhovia emeryi*. More specifically, adopting a pipeline similar to that in Cicconardi et al. (2017a, 2017b), the signatures of diversifying selection were searched in codon-based aligning groups of one-to-one orthologous ant genes with MACSE v. 1.01b (Ranwez et al. 2011), filtering with GBLOCKS v. 0.91b (Castresana 2000) under a “relaxed” condition (Parker et al. 2013; Cicconardi, Di Marino, et al. 2017; Cicconardi, Marcatili, et al. 2017) and using the aBSREL algorithm as implemented in the HYPHY batch language (Kosakovsky Pond et al. 2005) using a batch script (BRANCHSITEREL) in HYPHY (http://github.com/veg/hyphy).

In parallel, the same scOGs and HSP-OGs were scanned using the RELAX test (Wertheim et al. 2014) to search for putative signals of relaxation. In brief, RELAX tests the hypothesis of evolutionary-rate relaxation in selected branches of a phylogenetic tree compared with reference branches. A *k* value is computed to evaluate whether the selective strength *ω* shifts towards neutrality. The rate of d_N_/d_S_ (*ω*) can relax (*k* < 1), stay stable (*k* = 1), or intensify (*k* > 1). For both scOGs and HSP-OGs, and for both the ABSREL and RELAX tests, the Bonferroni-Holm sequential rejection procedure (Abdi 2010) was used to control the false discovery rate with a very stringent cutoff value of 0.005 for the adjusted *P-*values applied (Benjamin et al. 2017). Hsp60s and Hsp40s under relaxed selection were analyzed using the GENEMANIA prediction server (Warde-Farley et al. 2010) to predict their functions by creating an interaction network with genes by including protein and genetic interactions, pathways, co-expression, co-localization, and protein domain similarity.

Finally, the GOSTATS package for R (Falcon and Gentleman 2007) (settings: ANNOTATION org.Dm.eg.db; CONDITIONAL TRUE; TESTDIRECTION over) was used to check for GO terms of biological processes and KEGG pathway enrichments of scOGs under selection and relaxation using the whole set of tested scOGs as background. *P-*values < 0.005 and < 0.05 were implemented for GO terms and KEGG pathways, respectively.

### Synthesis of Hex-t1 gene and its enzyme activity assays

Following identification as under diversifying selection in *T. alpestre* (see results section “Dual signature of evolutionary pressures in single-copy OGs in the *T. alpestre* genome”), synthesis of genes and enzymes was attempted for phosphofructokinase, transketolase, phosphoglucosemutase, and Hex-t1, but it was successful only for the Hex-t1. In more detail, the Hex-t1 genes of *Tetramorium alpestre* (1398 bp) and *T. immigrans* (1386 bp) were synthesized by ThermoFisher lifetechnologies (Carlsbad, CA, USA) and Eurofins Genomics (Ebersberg, Germany), respectively. Both constructs carried a 5’ EcoRI and a 3’ HindIII recognition site for subsequent cloning as well as a C-terminal His-tag. The genes were excised from their carrier backbones by enzymatic digestion. The fragments were purified by agarose gel excision and sub-cloned into the pACEBac1 expression vector (MULTIBACTM, Geneva Biotech, Geneva, Switzerland). Correct insertion was verified by Sanger sequencing, and transformation into *D. melanogaster* cells and protein expression followed the standard protocols of the expression vector manufacturer. After cell harvesting, the presence of the recombinant protein in the cell lysates was verified by a western blot targeting the His-tag. The relative amounts of protein were quantified from the western blot using IMAGEJ v. 1.3 (https://imagej.net/Welcome). The cell lysates were centrifuged at 4000 rpm at 4 °C for 5 min, and the supernatant was directly used as protein source.

Enzyme activity assays followed a modified protocol of Crabtree & Newsholme (1972). Briefly, 1000 µl assay medium containing 75 mM Tris pH=7.5 (Merck Millipore, Burlington, MA, USA), 7.5 mM MgCl_2_ (Merck), 0.8 mM EDTA (Sigma-Aldrich, St. Louis, MO, USA), 1.5 mM KCl (Merck), 4.0 mM mercaptoethanol (Sigma), 0.4 mM NADP+ (Abcam, Cambridge, UK), 2.5 mM ATP (Abcam), and 10 mM creatine phosphate (Abcam) were placed in a disposable cuvette (Brand, Germany) in a SPECORD 210 PLUS UV/VIS spectrophotometer (Analytic Jena AG, Germany) with an attached 1157P programmable thermostate (VWR, Radnor, PA, USA). Variable amounts of D-glucose (VWR) and synthetic protein were added, and the cuvette was allowed to reach the assay temperature. Then, the reaction was started by adding 15 U creatine phosphokinase (Sigma) and 5 U glucose 6-phospahte dehydrogenase (Sigma).

The Hex-t1 activity was measured in triplicate as the rate of reduction of NADP^+^ causing an extinction at 340 nm. Each measurement lasted for 60 s, and every second, an extinction value was recorded. For the determination of the Michaelis-Menten constant (Km-value) of both enzymes, assays were conducted along a substrate gradient from 0.00167 to 24.30000 mM D-glucose, achieving saturated extinction curves. Assay temperatures ranged from 6 °C to 36 °C with 10 K increments. The software PRISM v. 8 (GraphPad Software, San Diego, CA, USA) was used to calculate the Km-values.

### Hex-t1 structure modelling and molecular dynamics simulations

The three-dimensional structures of *T. alpestre* and *T. immigrans* Hex-t1 proteins are not available. To study the effect of the amino acid changes on the proteins’ function/dynamics, we modeled the structures of these two proteins via homology modelling. The X-ray structure of the Hex-t1 isoform from *Schistosoma mansoni* (Mulichak et al. 2002), solved at 2.6 Å resolution was used as a template and the models were generated using the software MODELLER v. 9.21 (Webb and Sali 2014). Once we obtained the models, we built the simulative systems to run molecular dynamics simulations. The two proteins TalpHex-t1 and TimmHex-t1, in complex with the glucose substrate, were immersed in a triclinic box filled with TIP3P water molecules (Jorgensen et al. 1983) and rendered electroneutral by the addition of chloride counterions. The topology of the two systems, consisting of ∼ 80.000 atoms each, was built using the AMBER14 force field (Case et al. 2014), further converted in GROMACS v. 4.6 (Hess et al. 2008) format using ACPYPE v. 0.1.1 (Sousa Da Silva and Vranken 2012). Simulations were run on a GPU cluster using GROMACS v. 4.6 with the following protocol: 1) 25000 steps of steepest descent followed by 25000 steps of conjugate gradient minimization; 2) 5*100 ps of equilibration runs in constant volume and temperature (NVT) environment starting at 50 K and performed by increasing the temperature of 50 K after each run until a final value of 250 K; 3) 5*100 ps of equilibration runs in constant pressure and temperature (NPT) environment starting at 50 K and performed by increasing the temperature of 50 K after each run until a final value of 250 K; 4) the equilibrated systems were simulated for 500 ns at 279 or 300 K, i.e. at 6 or 26 °C, for a total of four simulations and 2 µs of sampling. Electrostatic interactions were considered by means of the Particle Mesh Ewald method (PME) with a cut off of 1.2 nm for the real space and Van der Waals interactions (Darden et al. 1993). Bond lengths and angles were constrained via the LINCS algorithm (Hess et al. 1997). The temperatures were kept constant at 279 or 300 ºK by using the velocity rescale with a coupling constant of 0.1 ps, and the pressure was kept at 1 bar using the Parrinello-Rahman barostat with a coupling constant of 1.0 ps during sampling (Parrinello and Rahman 1981). The four trajectories were collected and comparative analyses performed with the GROMACS suite or with in-house written code (available on request).

### Chill coma assay and quantitative qRT-PCR assay

The cold hardiness and expression of selected genes were compared between *T. alpestre* and *T. immigrans* using chill coma assays. From four colonies additional to those used for other experiments in this study, two per species, about 500 workers per colony were collected in August 2017; *T. alpestre*: Jaufenpass (18980: 46.83791° N, 11.29768° E) and Penser Joch (18978: 46.81402° N, 11.44198° E); *T. immigrans*: Vienna (18977: 48.12448° N, 16.43522° E.; 18983: 48.31341° N, 16.42529° E). Workers were collected alive using aspirators, transported to the laboratory in Innsbruck, and kept in polypropylene boxes (18.5 cm x 11 cm) with chambers of various sizes. The walls of the boxes were Fluon-coated (GP1, De Monchy International BV, Rotterdam, Netherlands) to prevent workers from escaping. Food (sugar-honey-water and deep-frozen *Drosophila* flies) and tap water were provided *ad libitum* three times a week. Workers were kept in a climate chamber (MIR-254, Panasonic, Etten Leur, Netherlands) at a 12L:12D photoperiod at constant 18 °C. The temperature of 18°C was the average of the monthly mean temperatures experienced at 200 m a.s.l. (*T. immigrans*, approx. 23.7 °C) and at 2000 m a.s.l. (*T. alpestre*, approx. 12.3 °C) in July 2017. Workers were acclimatized to 18 °C for at least three weeks before the chill coma assays.

After the acclimation period, from each of the four colonies, 90 workers were randomly chosen and divided into 18 equivalent pools of five workers, each of which were used for a single replicate of either the chill coma or the control assays. For the chill coma assay, worker pools were transferred into empty 5-ml glass vials, sealed, and immersed at 2300 hours Central European Summer Time (CEST) for 6.5 hours in a water:ethane-1,2-diol mix (1:1) bath set at −6 °C; in pilot experiments, this temperature had been determined as the highest temperature at which the ants had fallen into chill coma after 6.5 hours exposition; the temperature was identical for both species. Temperature was monitored using an electronic thermometer (TFX 430, ebro Electronic GmbH; Ingolstadt, Germany) with an accuracy of 0.05 °C inserted into an additional, empty vial treated in the same way as the vials with ants. After chill coma, at 0530 hours CEST, the vials were transferred to a climate chamber at 18 °C, and the workers were allowed to recover for 0.5, 2.0, and 4.0 hours, terminated at 0600, 0730, and 0930 hours CEST, respectively. For each recovery time, three replicate pools were used, and each chill-coma treatment and its corresponding control hence were replicated six-fold per species. After recovery, the workers were instantly transferred to RNAse-free reaction tubes and killed in liquid nitrogen. Tubes were stored at −70 °C. For the control assay, worker pools were transferred to empty glass vials and placed in the climate chamber at 18 °C. After 6.5 hours plus respective recovery time, the workers were killed and stored as described above.

For each recovery time and replicate, complete worker pools were used for the molecular analyses. RNA was extracted using the NUCLEOSPIN® RNA Kit (Macherey-Nagel, Düren, Germany) following the instructions of the manufacturer. First-strand cDNA was synthesized using 200 U REVERTAID reverse transcriptase (all reagents by Thermo Fisher Scientific, Waltham, USA), 40 U RIBOLOCK RNASE inhibitor, 5 µM random hexamer primers, 100 µM dNTPs, and 2 µl RNA extract in a total volume of 40 µl. The mixture was incubated for 5 min at 25 °C, 60 min at 42 °C, and 5 min at 70 °C on a UnoCycler 1200 (VWR, Radnor, USA). Quantitative PCR was conducted on a Rotorgene Q (Qiagen) PCR system. Each reaction contained 1× Rotor-Gene SYBR Green PCR Mastermix (Qiagen), 0.2 μM of primers, specifically designed for target genes (Table S12), and 1 µl cDNA in a total volume of 10 µl. All qRT-PCR reactions were performed as triplicates. *RpS20* was used as housekeeping gene as it had been established and used as housekeeping gene for comparable chill-coma assays in *Drosophila* (Colinet et al. 2010; Colinet et al. 2013). Cycling conditions were 95 °C for 5 min, followed by 40 cycles of 94 °C for 15 s, 58 °C for 10 s, and 72 °C for 15 s. Fluorescence was acquired at the end of each elongation step. PCR was followed by a melting curve analysis from 60 to 95 °C with 0.1 °C increments held for 5 s before fluorescence acquisition. Cycle threshold (Ct) values were subtracted from total number of cycles. On these values, linear-regression analyses using the software PAST v. 3.18 (Hammer et al. 2001) were based separately for each target gene, treatment, recovery time, and species (in each regression analysis, two populations and three replicate values were used, i.e., n = 6) using the housekeeping gene as independent variable and the target gene as dependent variable. For each target gene / recovery-time combination one-way analysis of covariance (ANCOVA) implemented in PAST followed by Bonferroni-Holm correction was then used to identify significant differences between treatments within species and between species within treatment. It is acknowledged that circadian rhythms in metabolic activities (Bloch et al. 2013) may be reflected in the data, in that the three different recovery times were initiated synchronously but terminated asynchronously. However, any such influence was controlled for as far as possible in the frame of this project in that each treatment with its corresponding control was synchronized as were the two species.

## Data access

The sequence data from this study have been submitted to the National Center for Biotechnology Information (NCBI) Sequence Read Archive (SRA) under BioProject numbers PRJNA532334, PRJNA533534, PRJNA533535, PRJNA533536 and PRJNA533537 (https://www.ncbi.nlm.nih.gov/bioproject/). The mtDNA genome assemblies have been submitted to the NCBI (GenBank accessions: MK861047 - MK861070). All accession codes of deposited and retrieved data are provided in Table S1.

## Acknowledgements

Philipp Andesner and Stefan Gross for valuable work in the laboratory. Research was supported by the Austrian Science Fund (FWF, P23409, P30861). The computational results presented here have been achieved in part by using the HPC infrastructure of the University of Innsbruck (LEO), the Vienna Scientific Cluster (VSC), the collaborative system of the Universities Innsbruck and Linz (MACH), and the Swiss National Supercomputing Centre (CSCS) hosted in Lugano.

