## Supplementary Figures for "Genomic signature of shifts in selection in a sub-alpine ant and its physiological adaptations"

***Supplementary Figure 1:*** *K-mer distribu ons of ant assemblies.............................. pg 2*

***Supplementary Figure 2:*** *GC content distributions across the Tetramorium ant genome assemblies.......................................................................................................... pg 3*

***Supplementary Figure 3:*** *Tetramorium ssp. GC content versus sequencing depth. pg 4*

***Supplementary Figure 4:*** *Coverage distributions..................................................... pg 5*

***Supplementary Figure 5:*** *Ant repeat versus assembly size....................................... pg 6*

***Supplementary Figure 6:*** *Estimate genome assembly completeness in ants.......... pg 7*

***Supplementary Figure 7:*** *Gene feature distributions................................................ pg 8*

***Supplementary Figure 8:*** *Gene similarity and completeness.................................. pg 9*

***Supplementary Figure 9:*** *MtDNA gene synteny in ants.......................................... pg 10*

***Supplementary Figure 10:*** *Ant nuDNA phylogeny tree........................................... pg 11*

***Supplementary Figure 11:*** *nuDNA MP-EST phylogenetic tree of ants.................. pg 12*

***Supplementary Figure 12:*** *Dated nuDNA phylogeny of ants................................. pg 13*

***Supplementary Figure 13:*** *Dated mtDNA phylogeny of ants................................. pg 14*

***Supplementary Figure 14:*** *Phylogeny of Hsp90s in ants....................................... pg 15*

***Supplementary Figure 15:*** *Phylogeny of Hsp70s in ants....................................... pg 16*

***Supplementary Figure 16:*** *Phylogeny of Hsp60s in ants....................................... pg 17*

***Supplementary Figure 17:*** *Phylogeny of Hsp40s in ants....................................... pg 18*

***Supplementary Figure 18:*** *Phylogeny of sHsps in ants.......................................... pg 19*

***Supplementary Figure 19:*** *Function prediction by interacting networks Function prediction by interacting networks.......................................................................... pg 20*

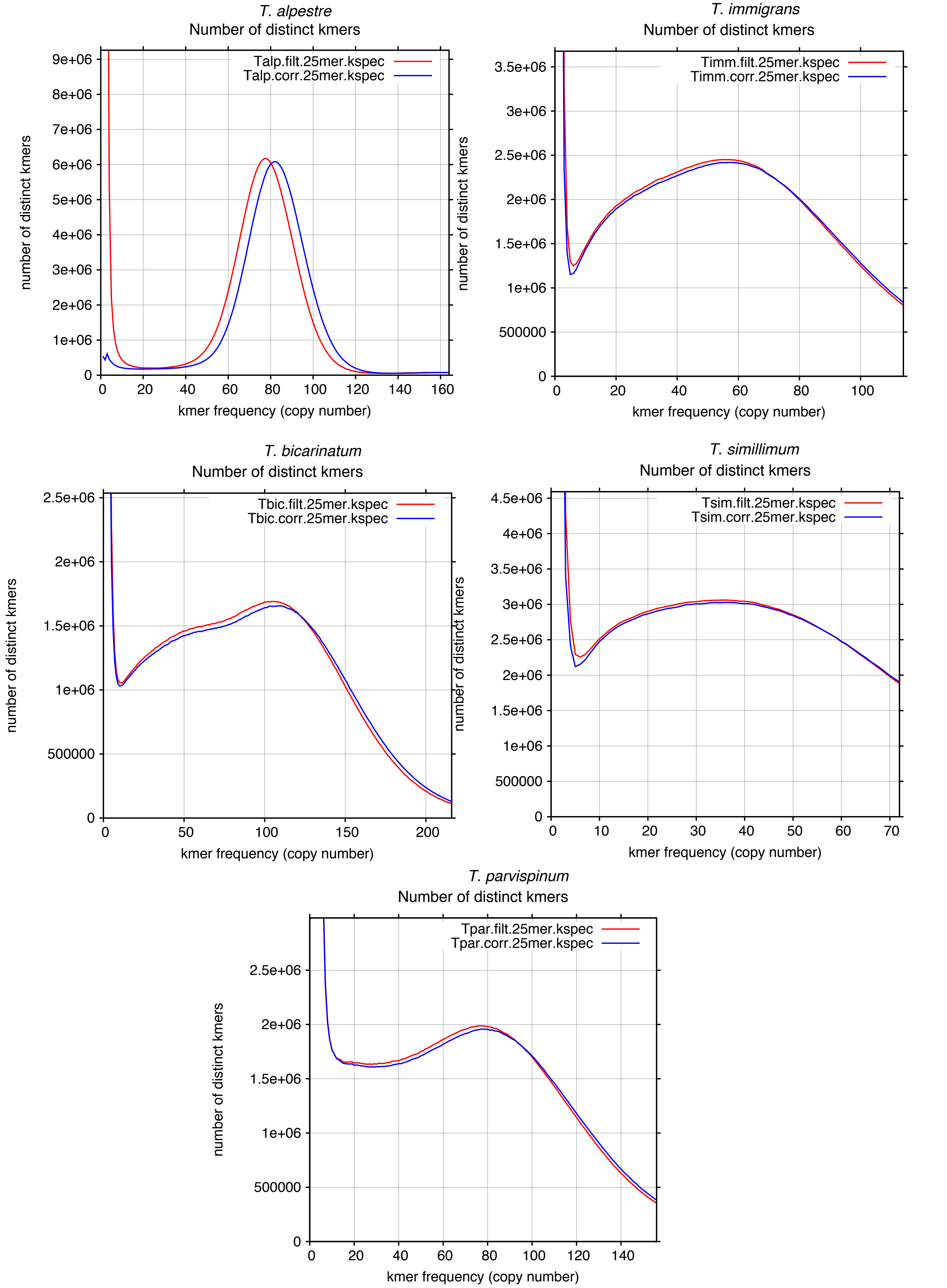


**Supplementary Figure 1.** ***K*-mer distributions of ant assemblies**
25-mer frequency distributions of *Tetramorium* ant genome assemblies, using all raw and error-corrected reads for each assembly.

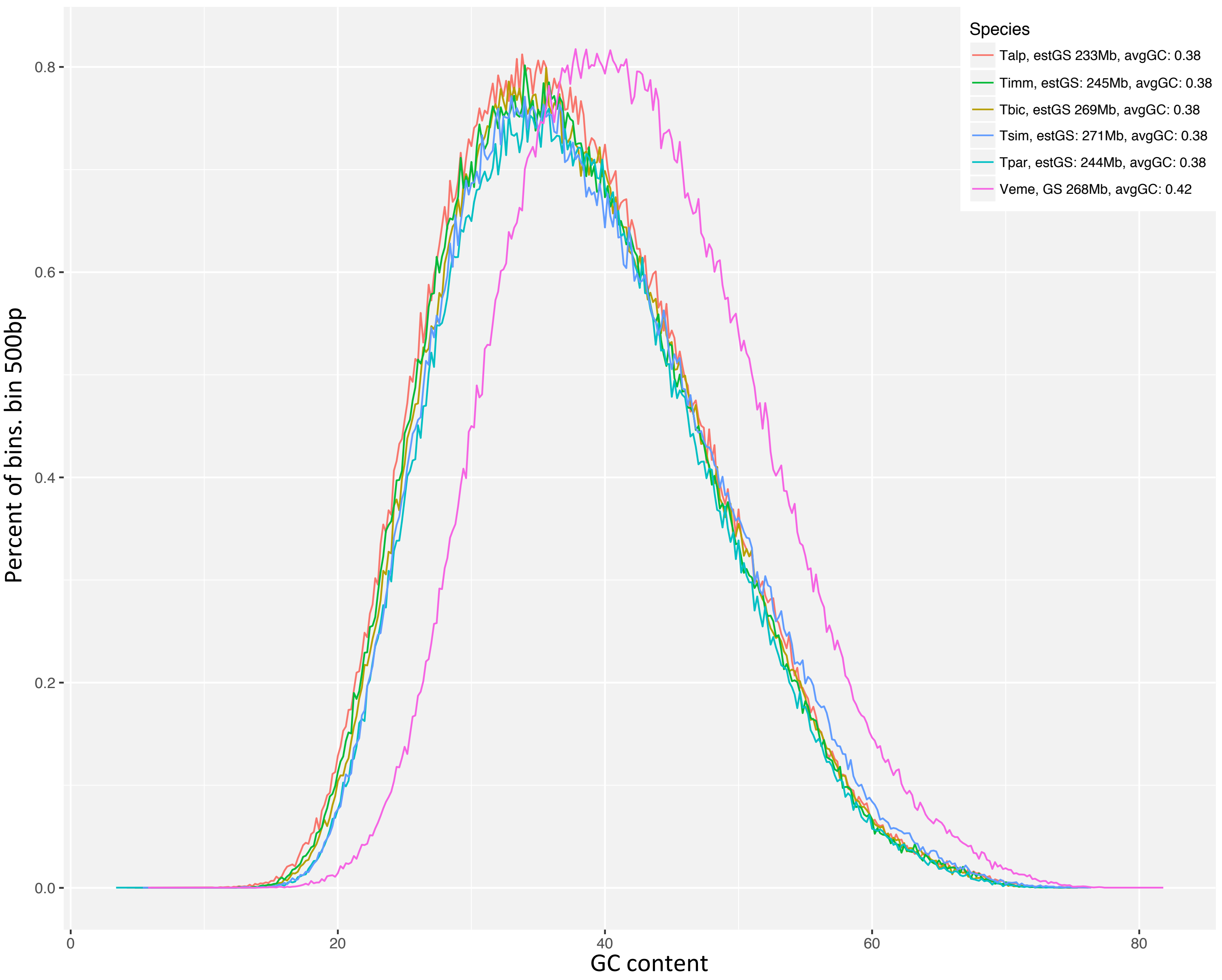


**Supplementary Figure 2. GC content distributions across the *Tetramorium* ant genome assemblies**
The *x*-axis shows the GC content and the *y*-axis the proportion of non-overlapping sliding windows of 500 bp. The legend shows species IDs (*T. alpestre* (Talp), *T. immigrans* (Timm), *T. bicarinatus* (Tbic), *T. simillimum* (Tsim), *T. parvispinum* (Tpar), *V. emeryi* (Veme), genome assembly sizes, and the average GC content. The ant genome assemblies have very similar GC content distributions with a peak of approximately 30% GC.

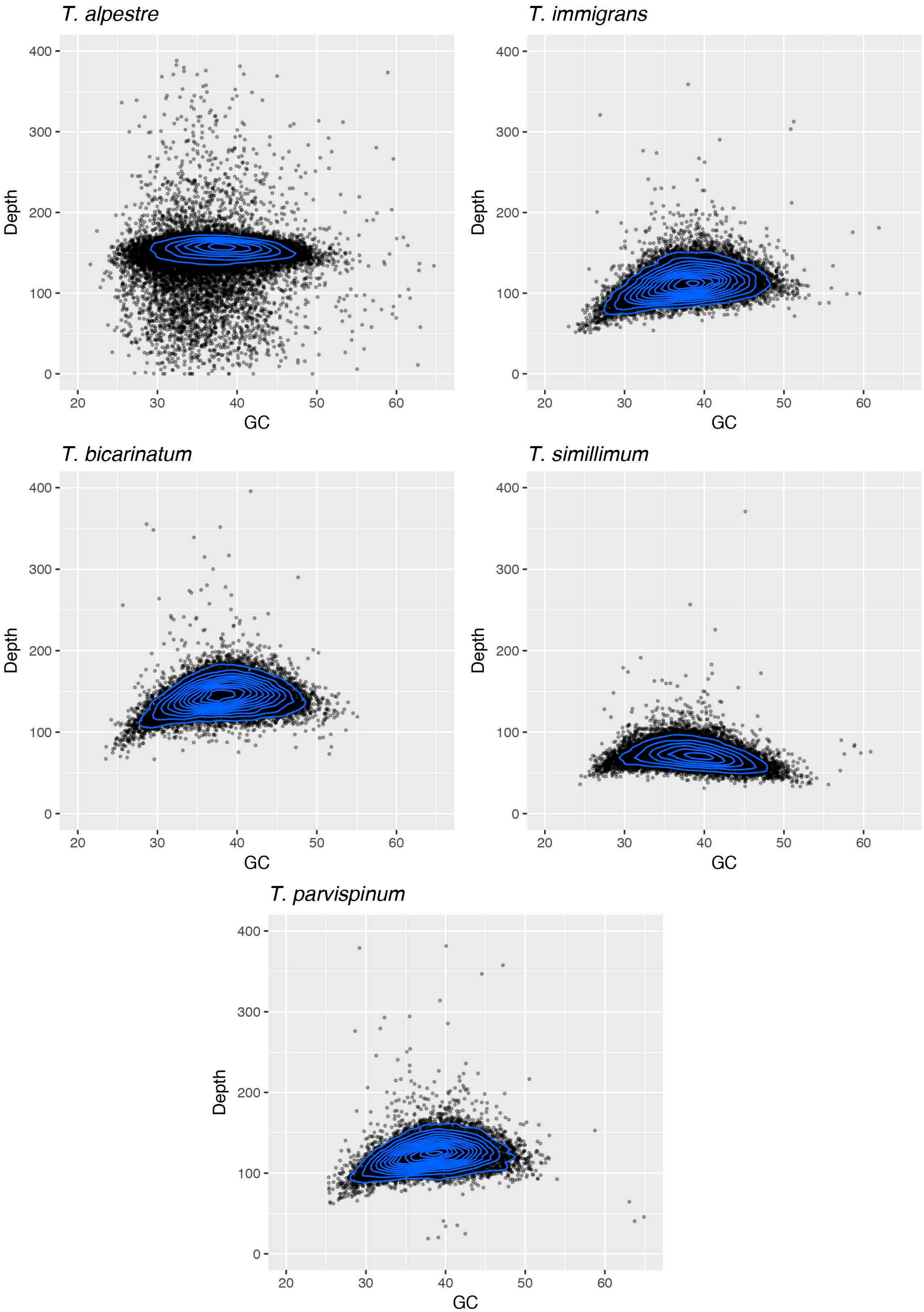


**Supplementary Figure 4.** ***Tetramorium ssp.*** **GC content versus sequencing depth**

Density plot showing the correlation between GC content and sequencing depth in the five Tetramorium ant genome assemblies. The *x*-axis represents GC content; the *y*-axis represents average sequencing depth, in a on-overlapping sliding windows 10 kb.

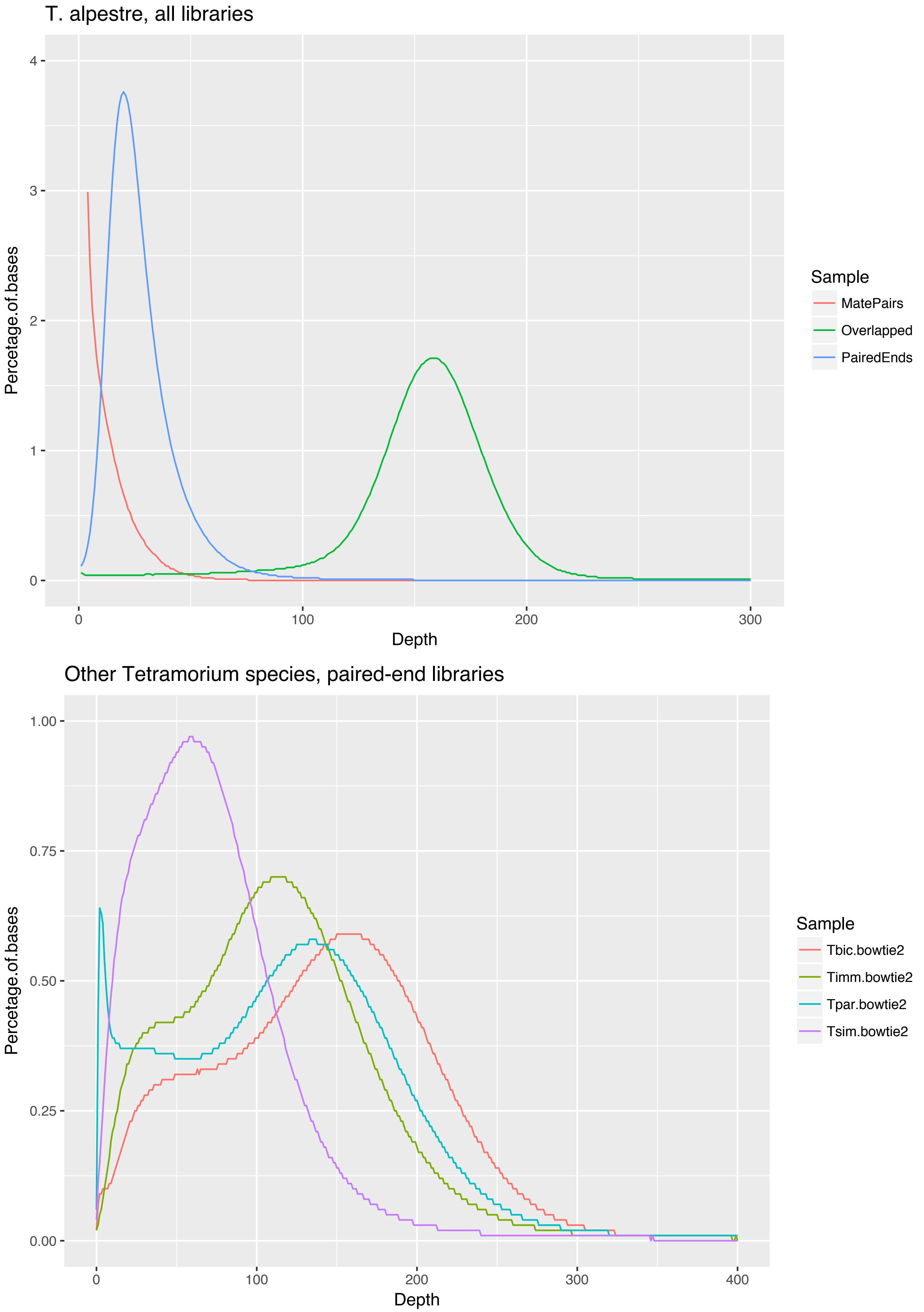


**Supplementary Figure 4. *Tetramorium* coverage distribution**

Density plot showing depth of coverage distributions for the Tetramorium ant genome assemblies. The *x*-axis represents the depth of coverage, while the *y*-axis the proportion of total bases at a given depth (abbreviations as above).

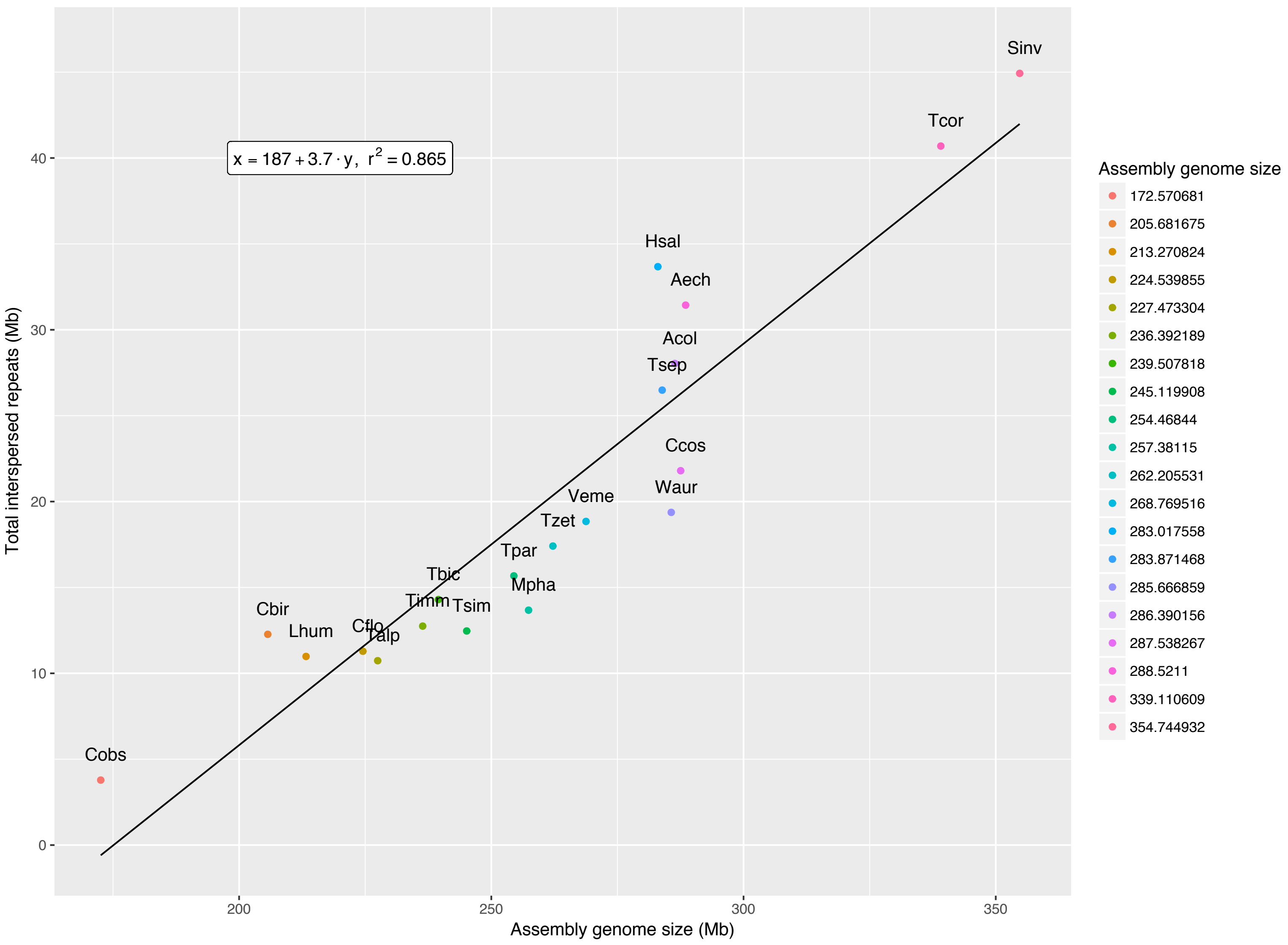


### Supplementary Figure 5. Ant repeat versus assembly size

Scatter plot correlating assembly size in ants (*x*-axis) versus total interspersed repeat content (*y*-axis) in 20 ant genome assemblies. These numbers are linearly correlated (Pearson rho = 0.976, *P* value < 0.005), with an intercept around 187 Mb (abbreviations as above, plus *Solenopsis invicta* (Sinv),  *Monomorium pharaonis* (Mpha), *Wasmannia auropunctata* (Waur), *Acromyrmex echinatior* (Aech), Trachymyrmex cornetzi (Tcor), Trachymyrmex zeteki (Tzet), Cyphomyrmex costatus (Ccos), *Camponotus floridanus* (Cflo), *Linepithema humile* (Lhum), *Cerapachys biroi* (Cbir), *Harpegnathos saltator* (Hsal), *Lasius niger* (Lnig), *Pogonomyrmex barbatus* (Pbar), *Atta cephalotes* (Acep)).

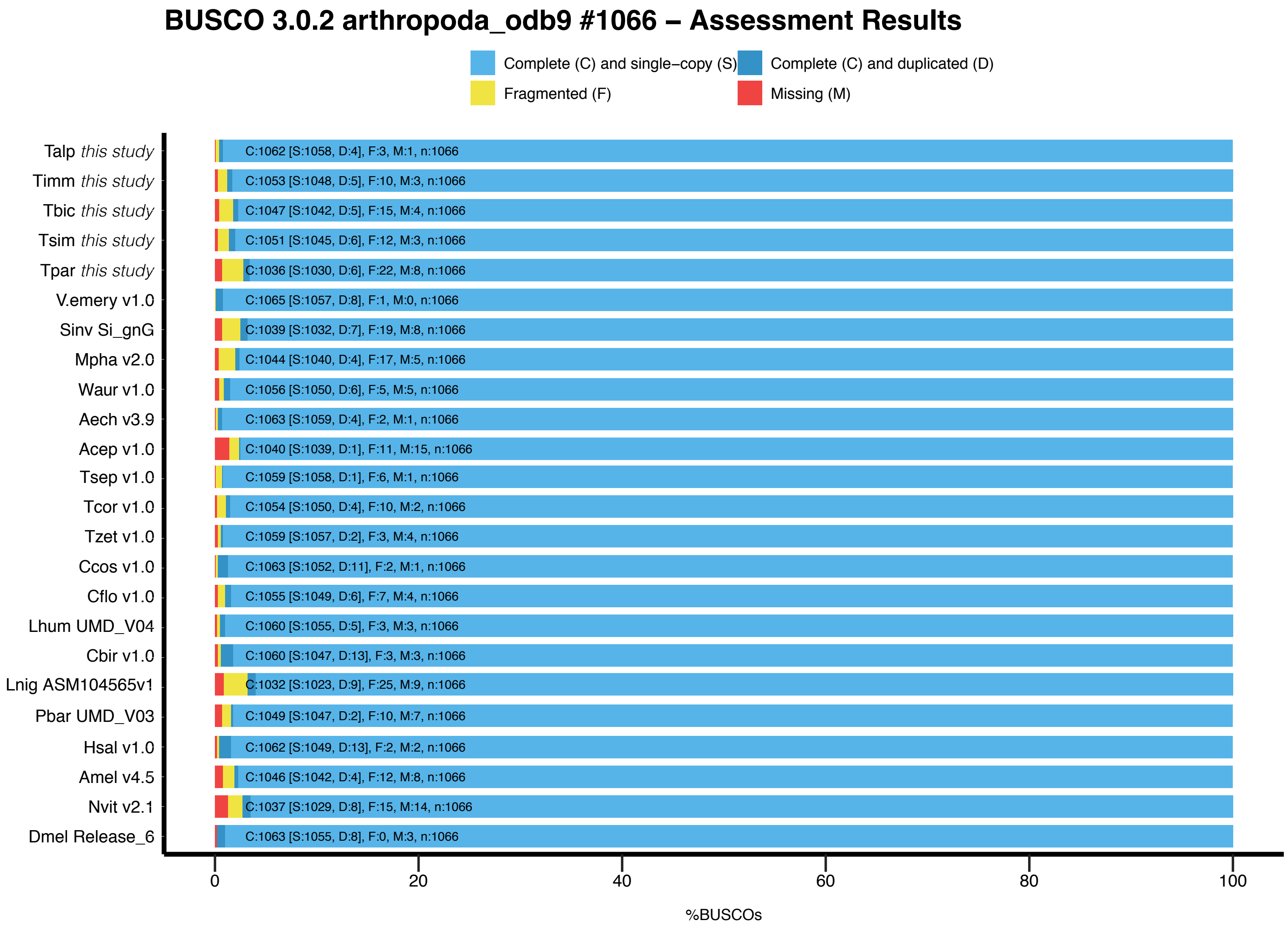


**Supplementary Figure 6. Estimate genome assembly completeness in ants**

Stacked histogram of Universal Single-Copy Orthologs (BUSCO) in five assembled *Tetramorium* species compared with the available ant genome assemblies, and three out groups species (abbreviations as above, plus *Nasonia vitripennis* (Nvit), *Apis mellifera* (Amel) and *Drosophila melanogaster* (Dmel)).

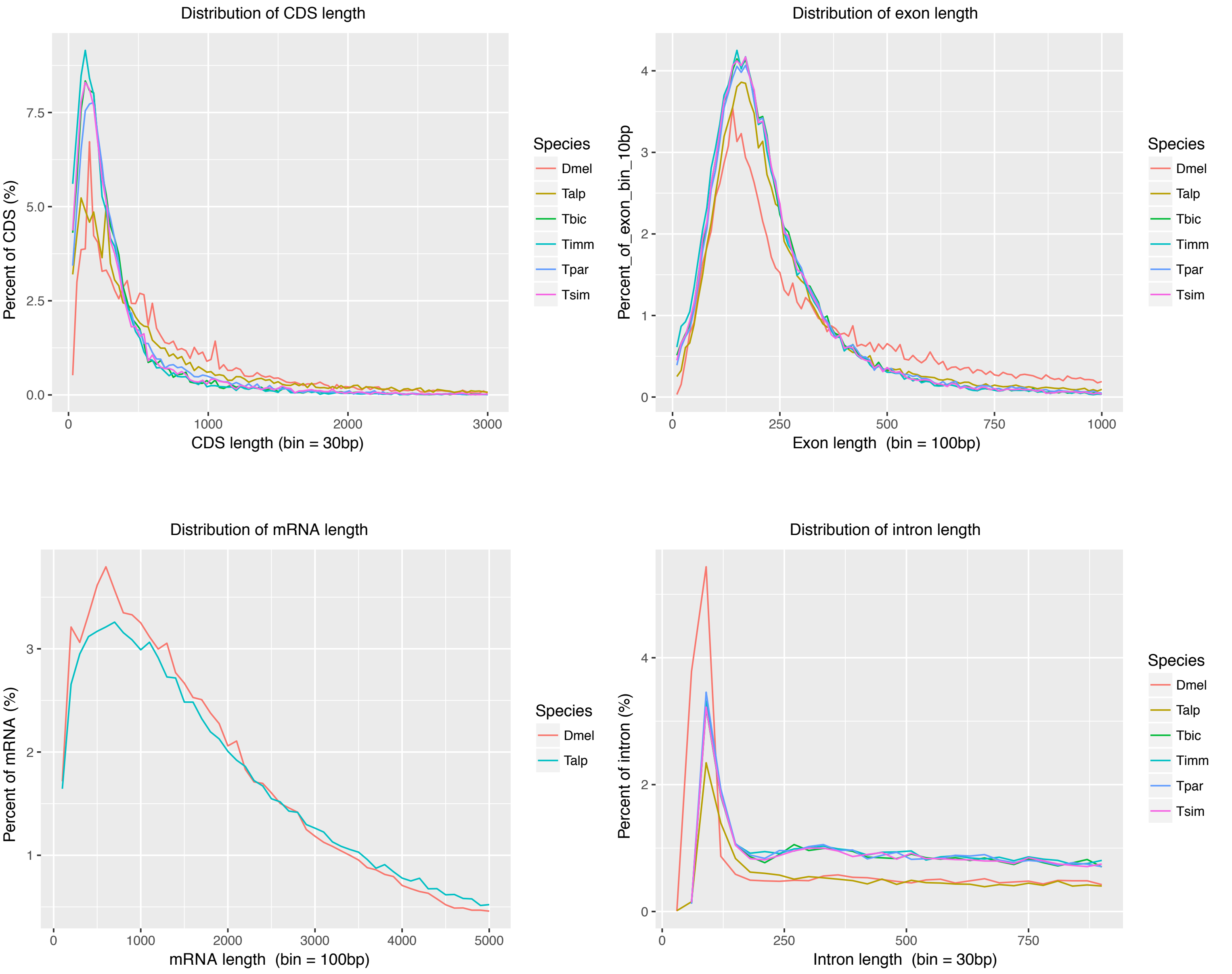


**Supplementary Figure 7. Gene feature distributions**

Length distributions for four general features of the final *Tetramorium* ant gene annotation sets. The corresponding distributions for *Drosophila melanogaster* (Dmel) is included for comparison.

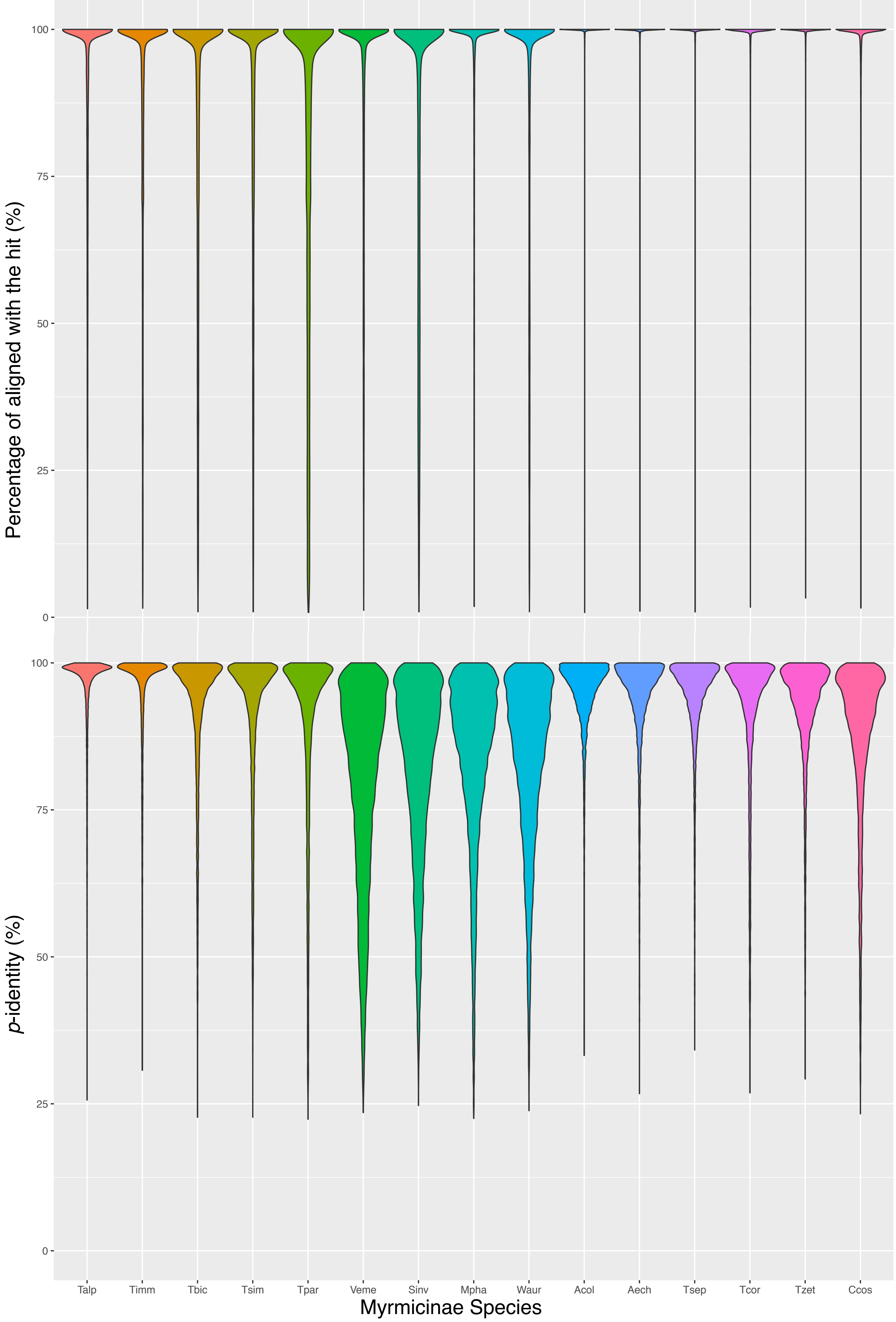


**Supplementary Figure 8. Gene similarity and completeness**

Violin plot showing the degree of gene completeness in terms of fraction (upper panel), and the *p*-identity distributions of aligned gene (lower panel) within all ant transcriptomes.

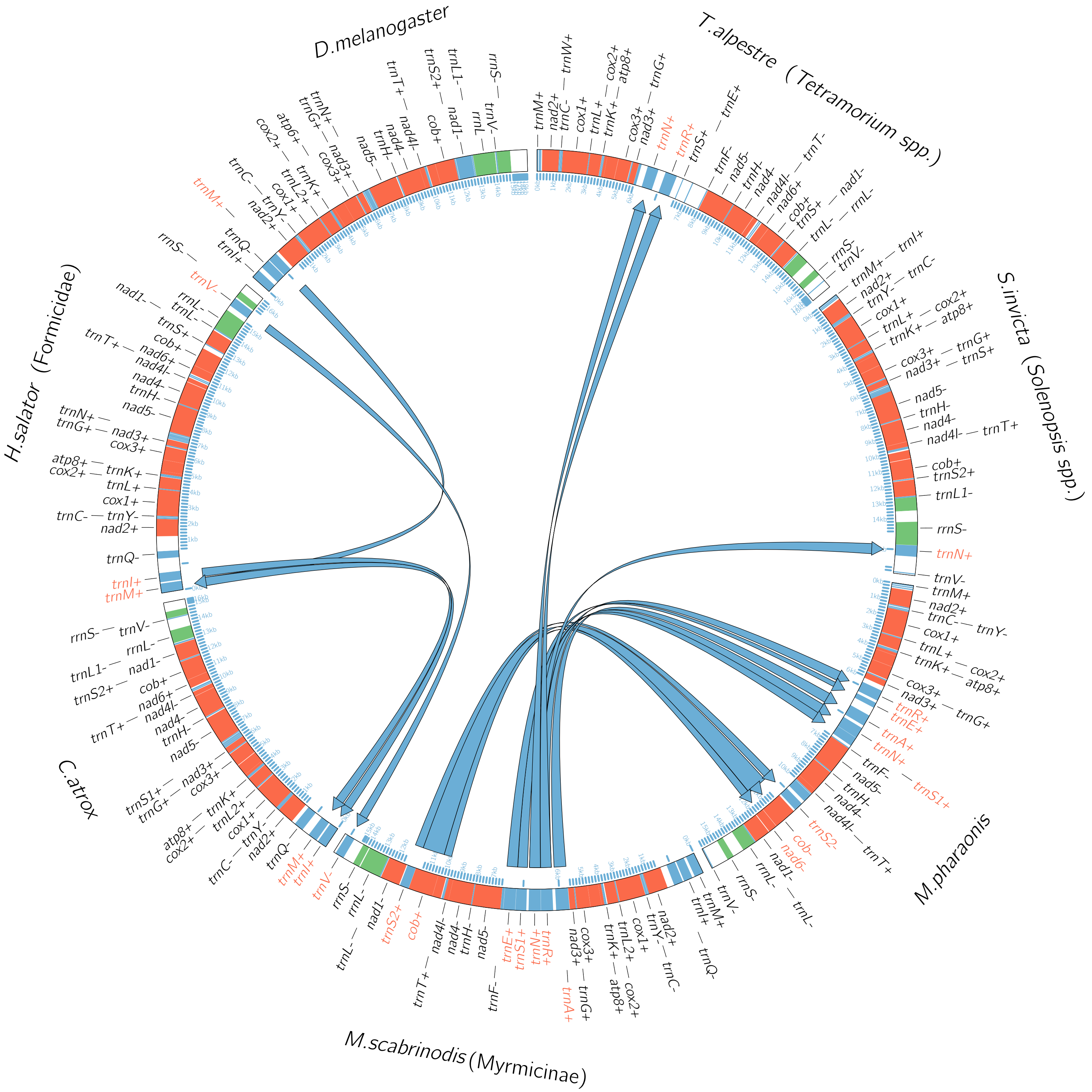


**Supplementary Figure 9. MtDNA gene synteny in ants**

Diagram showing the synteny and genetic rearrangments of mtDNA across the 39 studied ants. *Drosophila melanogaster* was considered as the ancestral state, and all the other species are listed in a phylogenetic order, in anticlockwise sense. The gene order in Harpegnathos saltator is considered as the most basal stat from which all Formicide are derived from. *Camponotus atrox* has its specific rearrangments, *Myrmica scabrinodis* instead is derived derived from the *H. saltator* arrangment and is considered the basal gene order from where *Monomorium pharaonis*, *Solenopsis spp.*, and *Tetramorium spp.* are derived from. Genes subject to rearrangment are listed in red, and region magnified.

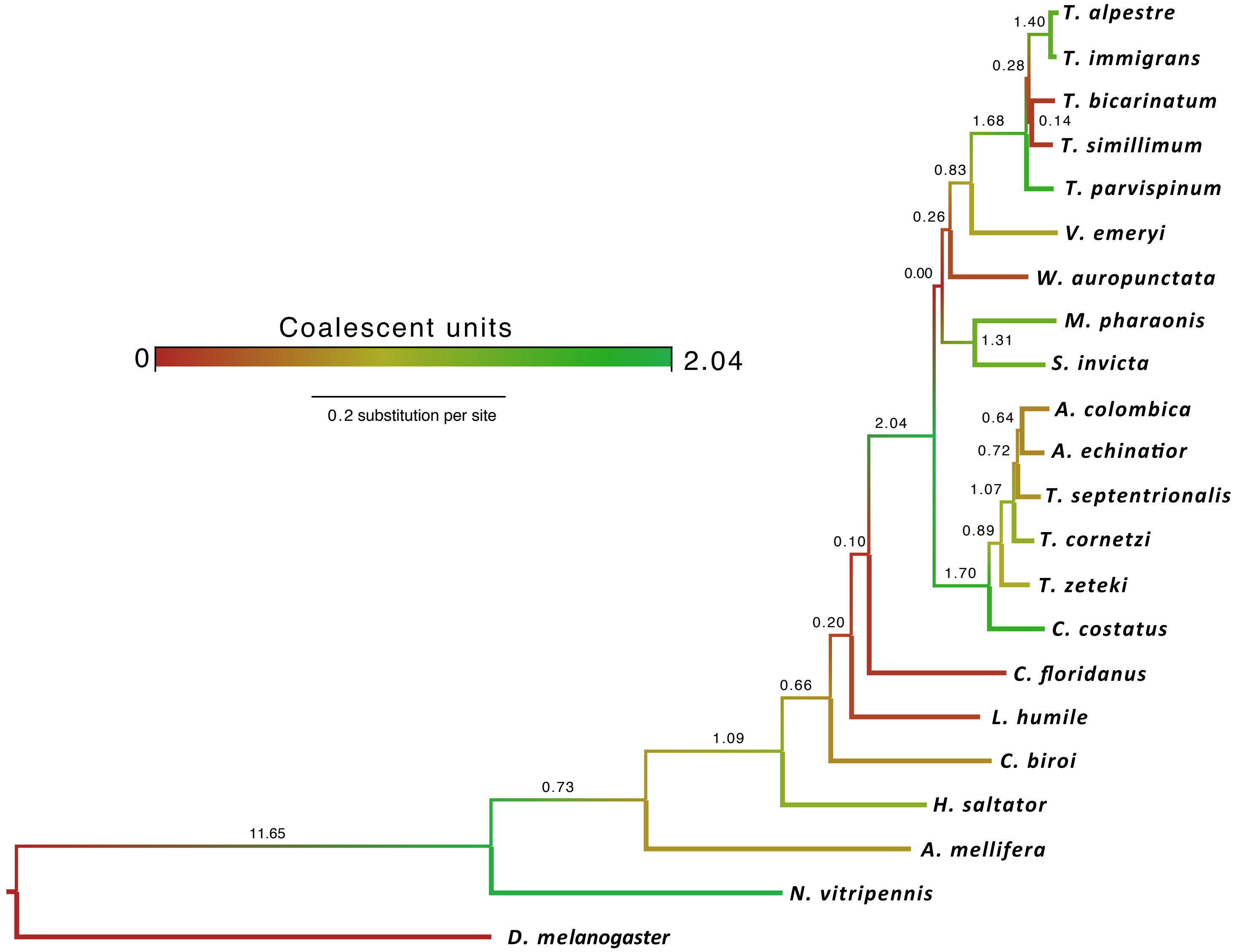


**Supplementary Figure 10. Ant nuDNA phylogeny tree**

The most likely tree resulting from a maximum-likelihood analysis of 8.5 Mb sites (3261 concatenated scOGs) and 19 ant species and the *D. melanogaster*, *N. vitripennis* and *A. mellifera* as outgroup. All nodes were supported by bootstrap frequencies. Single gene trees were also used to compute an no-concatenated tree. Colour and numbers on branches indicate the overall coalescent units (CU).

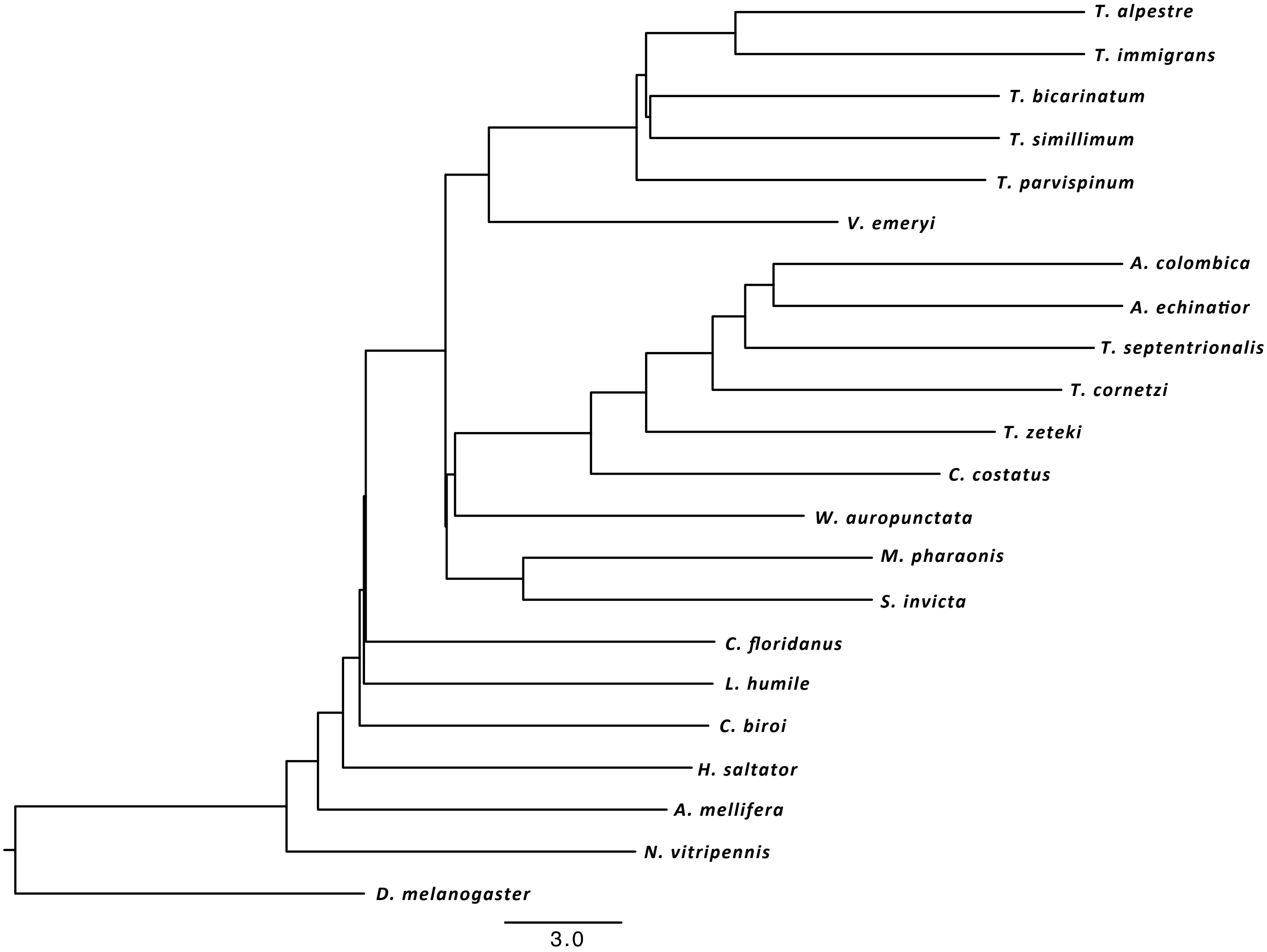


**Supplementary Figure 11. nuDNA MP-EST phylogenetic tree of ants**

Phylogenetic tree for 19 ant species plus three outgroups inferred summarising 3261 scOG gene trees with MP-EST, using the STRAW web-server. Branch lengths represent coalescent units (CU).

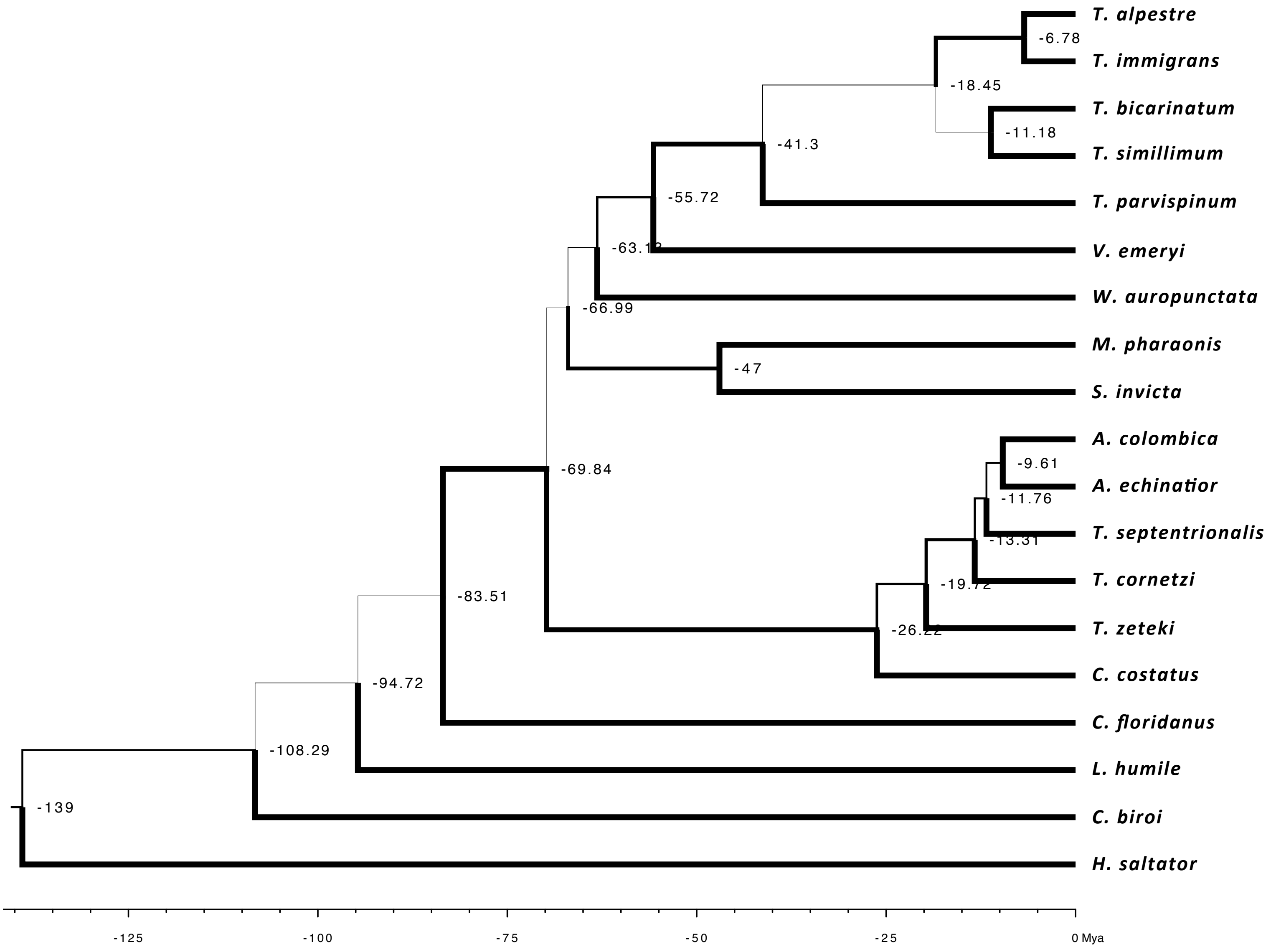


**Supplementary Figure 12. Dated nuDNA phylogeny of ants**

A dated time tree for 19 ant species inferred after using a penalized likelihood approach on the nuDNA phylogeny. Branch thickess is proportional to the CF and numbers on nodes indicate dates in millions of years before present.

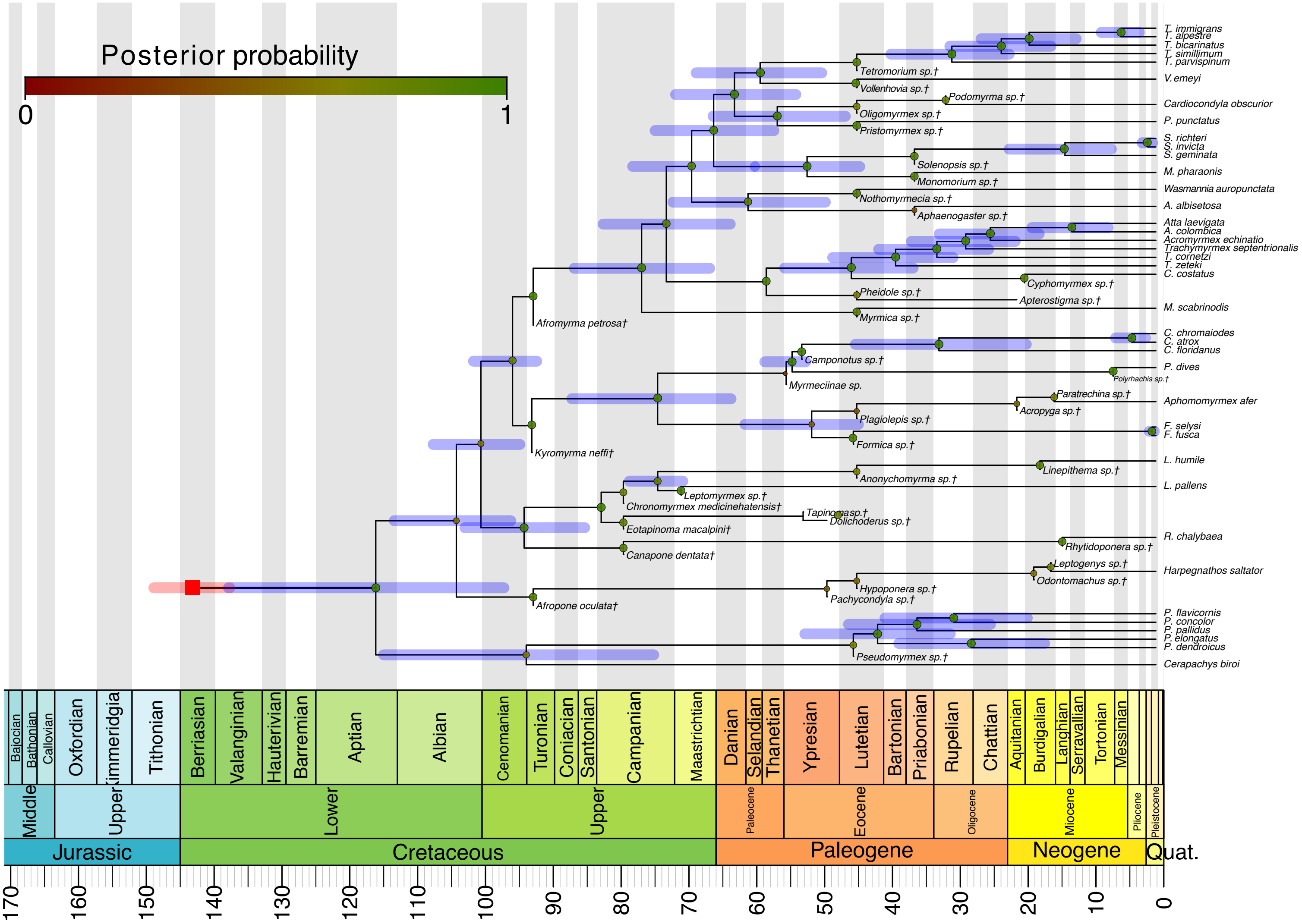


**Supplementary Figure 13. Dated mtDNA phylogeny of ants**

The maximum clade credibility tree summarized by TreeAnnotator and plotted with a geological timescale using the strap package in R. The fossil taxa are all indicated with an † in the taxon names. The red square represents the mean origin time. The remaining internal nodes of the tree are indicated with circles, colours mark posterior probability. The 95% credible intervals for node ages are shown with transparent bars, for only nodes that are represented in the extant tree.

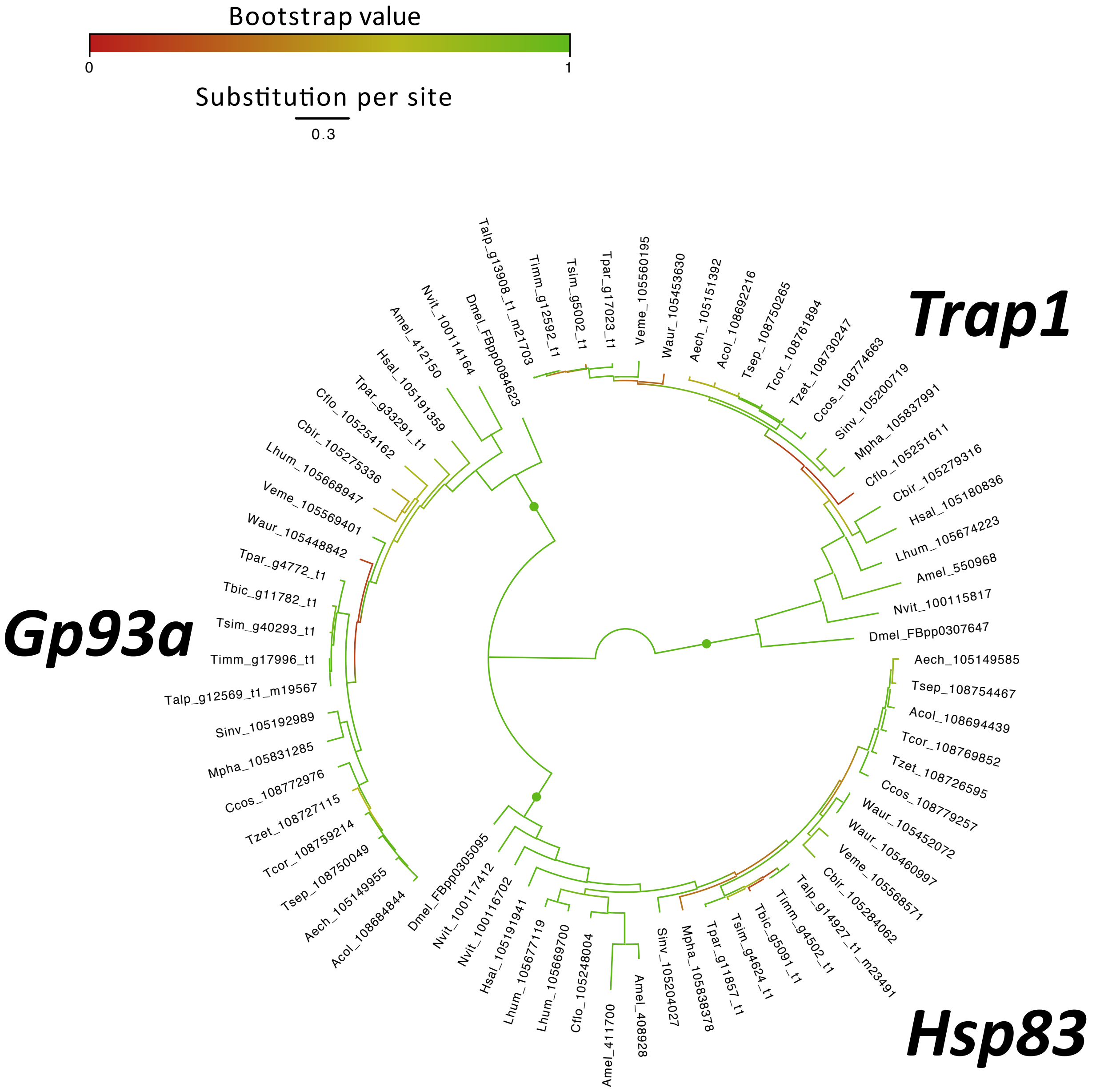


**Supplementary Figure 14. Phylogeny of Hsp90s in ants**

The maximum-likelihood phylogeny of Hsp90s in ants using nucleotide sequences. In color from red to green is represented the bootstrap values of each node. Green circles correnspond to orthologous gene clusters.

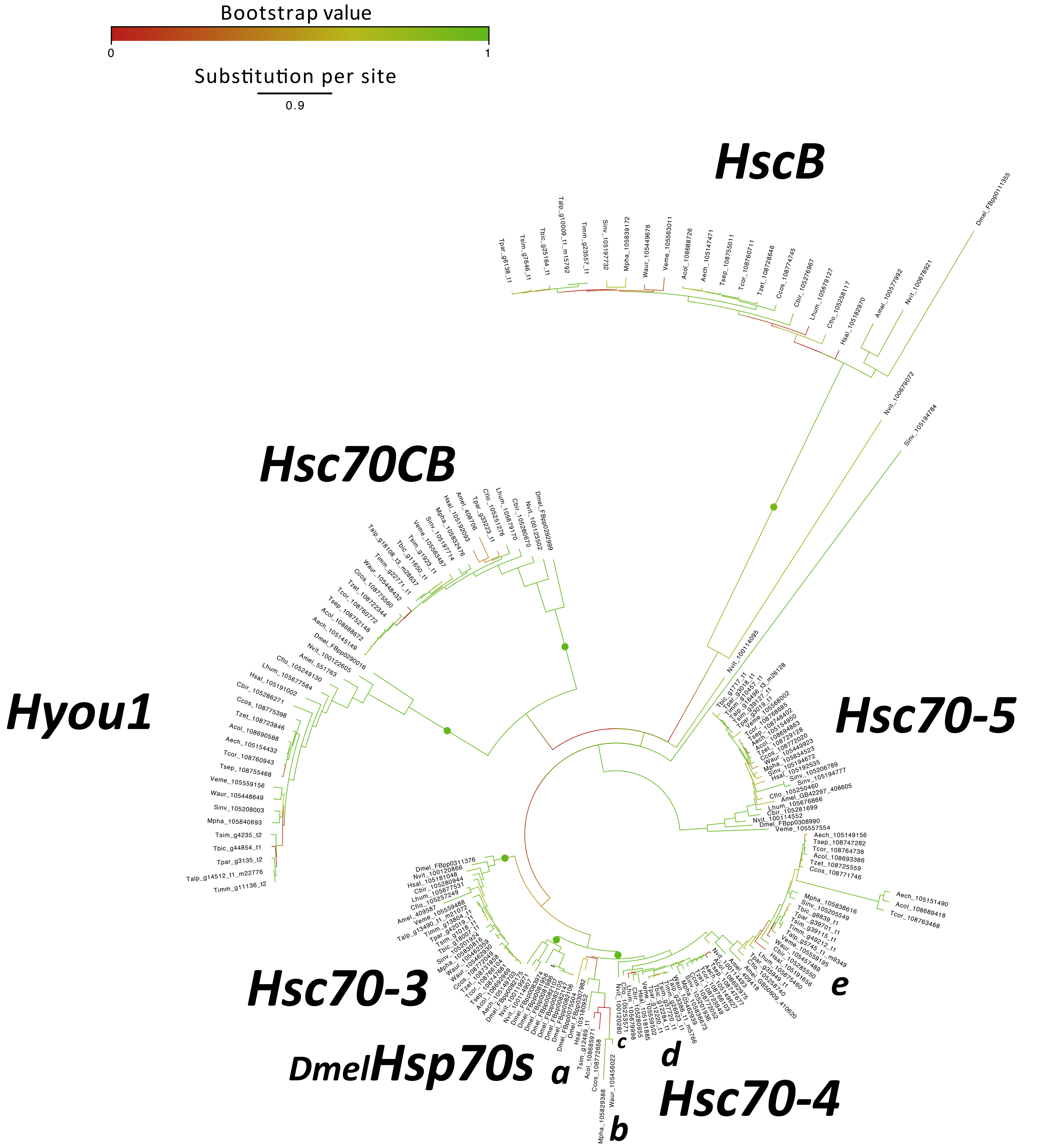


**Supplementary Figure 15. Phylogeny of Hsp70s in ants**

The maximum-likelihood phylogeny of Hsp70s in ants using nucleotide sequences. In color from red to green is represented the bootstrap values oof each node. Green circles correnspond to orthologous gene clusters.

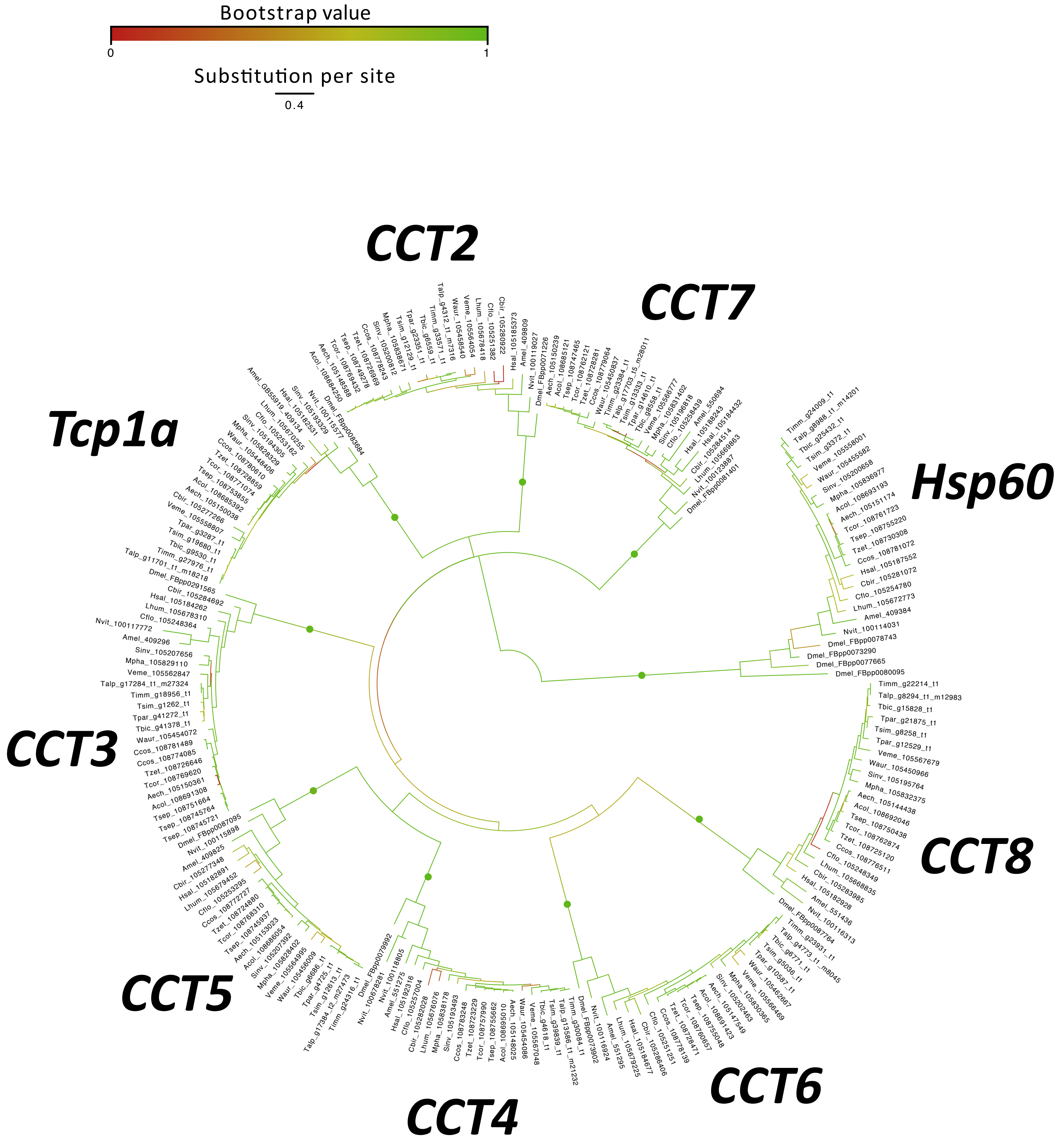


**Supplementary Figure 16. Phylogeny of Hsp60s in ants**

The maximum-likelihood phylogeny of Hsp60s in ants using nucleotide sequences. In color from red to green is represented the bootstrap values oof each node. Green circles correnspond to orthologous gene clusters.

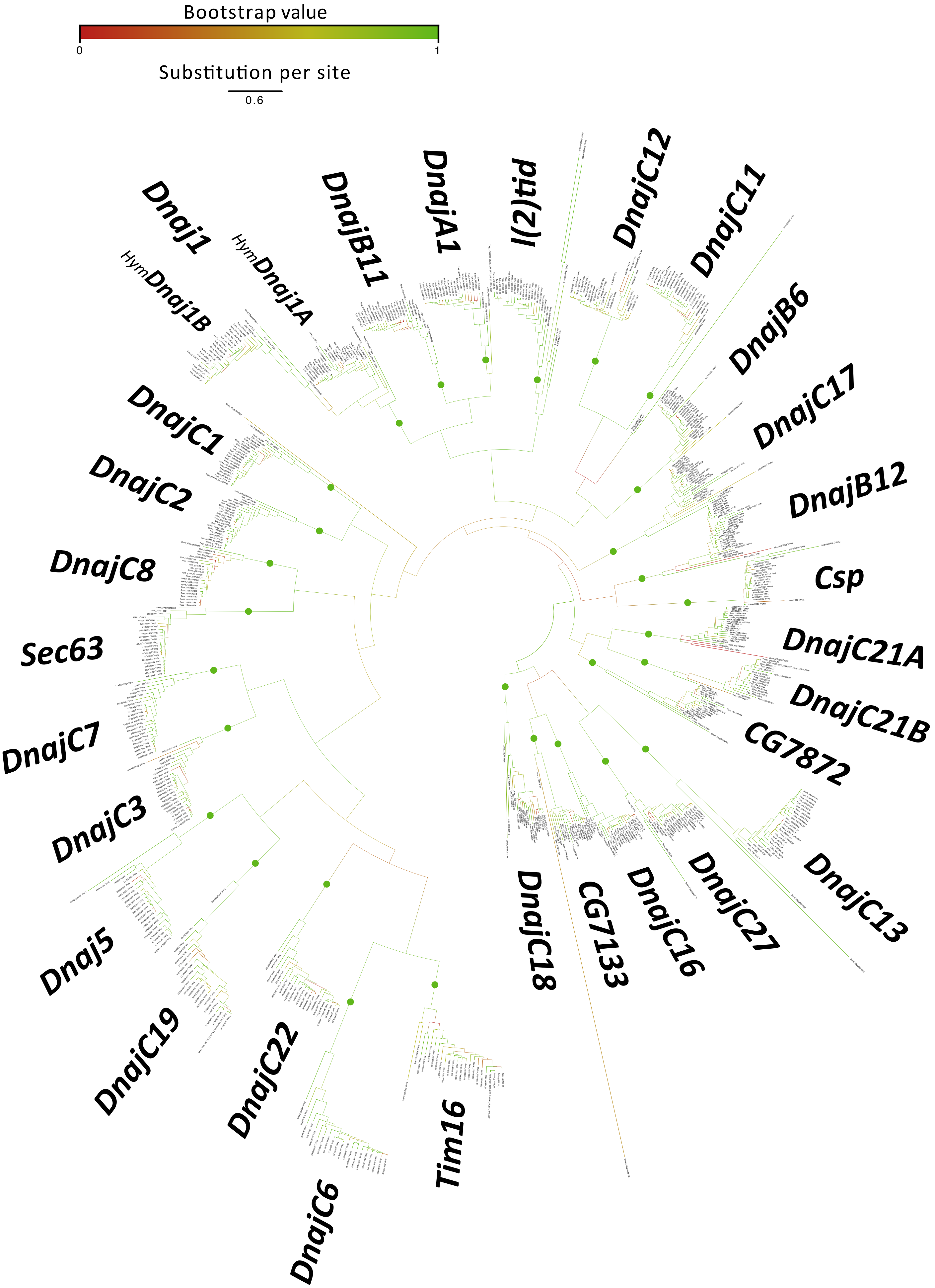


**Supplementary Figure 17. Phylogeny of Hsp40s in ants**

The maximum-likelihood phylogeny of Hsp40s in ants using nucleotide sequences. In color from red to green is represented the bootstrap values oof each node. Green circles correnspond to orthologous gene clusters.

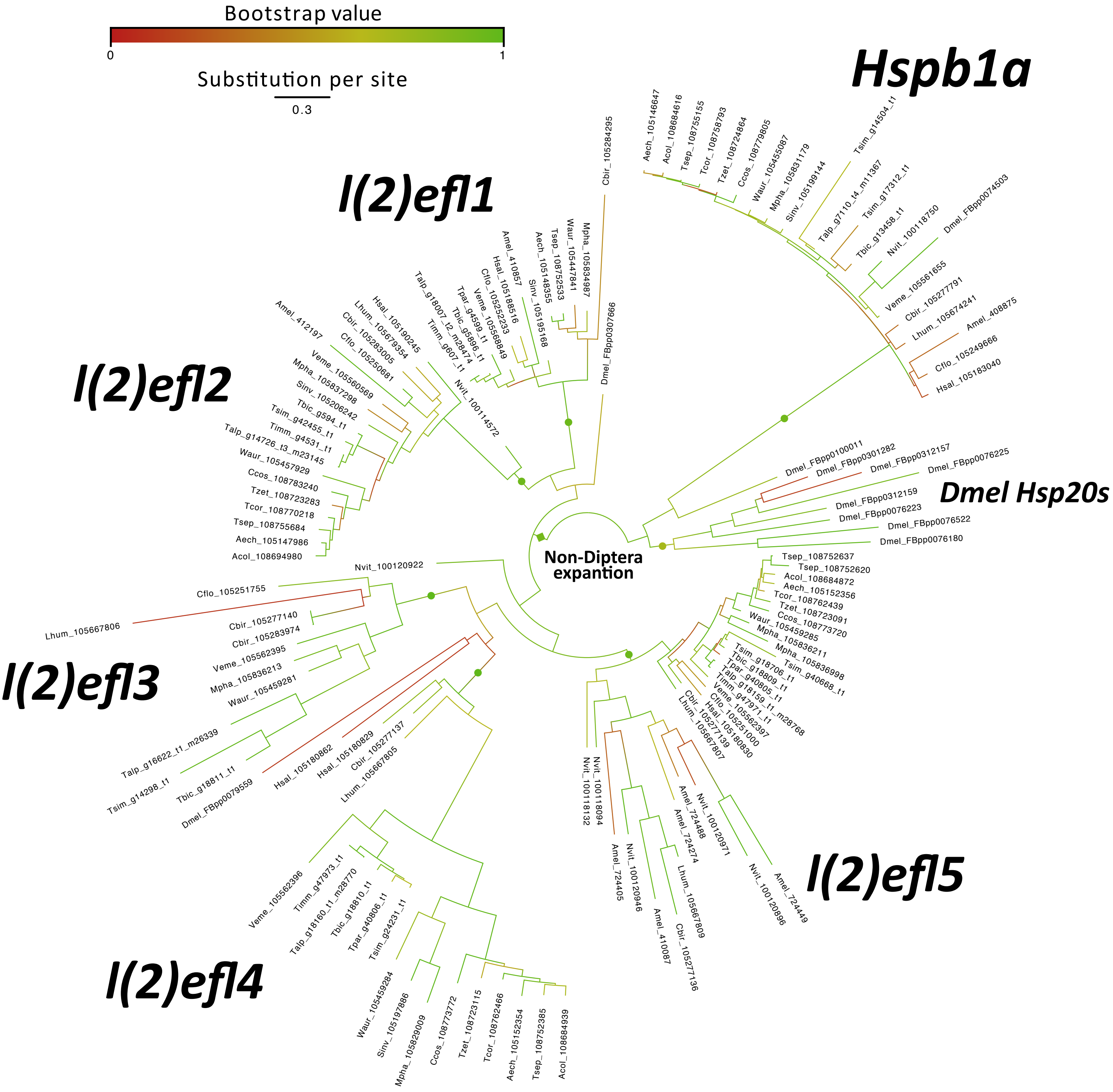


**Supplementary Figure 18. Phylogeny of sHsps in ants**

The maximum-likelihood phylogeny of sHsps in ants using nucleotide sequences. In color from red to green is represented the bootstrap values oof of each node. Green circles corrensponds to orthologous gene clusters.

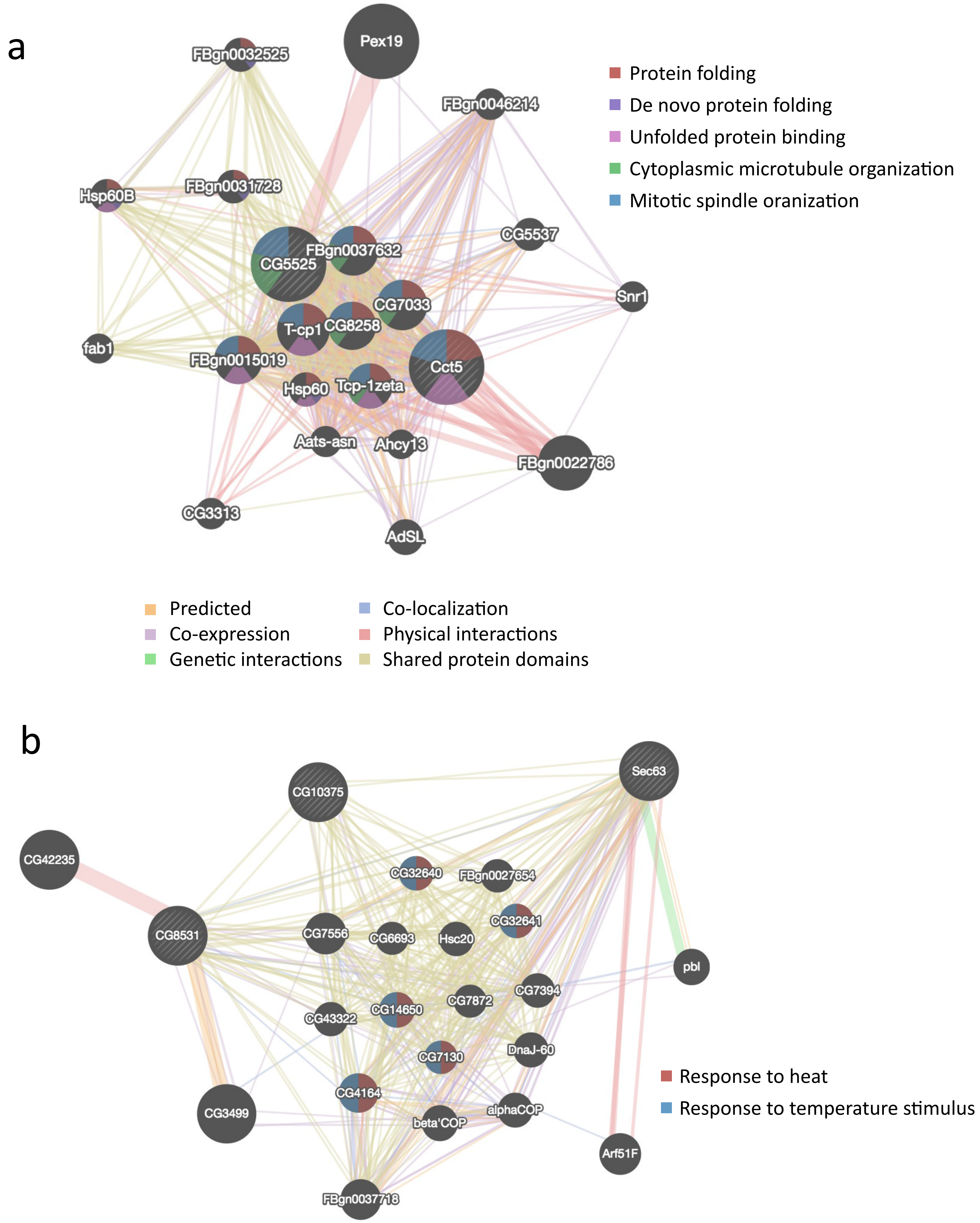


**Supplementary Figure 19. Function prediction by interacting networks**
Composite network of protein interaction for a) Hsp60s: Cct4 (CG5525) and *Cct5*; and b) Hsp40: *Sec63*, *DnajC8* (CG10375), DnajC11 (CG8531). The nodes are genes and the edges are the type of relationship, the thickness of edges represent the amount of evidences supporting the relationship.
